# *Rsc3K,* a SMV resistance gene localized on chromosome 2 in soybean cv. Kefeng No.1, interacts with virulence determinant P3

**DOI:** 10.1101/2021.12.30.474612

**Authors:** Tongtong Jin, Jinlong Yin, Song Xue, Bowen Li, Tingxuan Zong, Yunhua Yang, Hui Liu, Mengzhuo Liu, Kai Xu, Kai Li, Liqun Wang, Haijian Zhi

## Abstract

*Soybean mosaic virus* (SMV) is one of the most devastating viral pathogens in *Glycine max* (L.) Merr (soybean). Twenty-two SMV strains (SC1-SC22) isolated in China were identified based on their responses to ten soybean cultivars. By using the F_2_-derived F_3_ (F_2:3_) and recombinant inbred line (RIL) populations of resistant Soybean cultivar (cv.) Kefeng No.1 × susceptible cv. Nannong 1138-2, we localized the gene mediating resistant to SMV-SC3 strain to a 90 kb interval on chromosome 2 in Kefeng No.1. Bean pod mottle virus (BPMV)-induced gene silencing (VIGS) were used to study the gene function of candidate genes in the mapping interval and revealed that an recombinant gene, later named as *Rsc3K*, caused by internal deletion of a genomic DNA frag ement in Kefeng No.1, is the resistant gene to SMV-SC3. By shuffling genes between avirulent isolate SC3 and avirulent SMV isolate 1129, we found that P3 is the virulence determinant causing resistance on Kefeng No.1. We showed the interaction between Rsc3K and P3 by the yeast two-hybrid (Y2H) and bimolecular fluorescent complementation (BiFC) assays. In conclusion, this study demonstrated that *Rsc3K* plays a crucial role in resistance of Kefeng No.1 to SMV-SC3 by direct interaction with viral protein P3.

**Highlight:** Rsc3K resistant to SMV-SC3 by interacting with P3.

## INTRODUCTION

Soybean is the most important legume crop in the world due to its high protein and oil contents. *Soybean mosaic virus* (SMV) is a severe threat to soybean production in China as well as in the rest of the world (Zhang *et al*., 1999). In soybean-producing areas, the annual yield loss due to soybean mosaic virus disease is 5 to 7% (Zhang 2009), yield of severe diseased soybean field can drop 50-80% (Mohammad and Sher 2002).

In order to systematically study soybean mosaic virus, SMV isolates have been classified into different strains according to the pathogenic phenotype of SMV in different soybean cultivars. In the United States, Cho and Goodman used 8 soybean cultivars to classify 98 SMV isolates into seven strains, namely, G1 to G7 (Cho and Goodman 1979, 1982). Four dominant genes, *Rsv1*, *Rsv3*, *Rsv4,* and *Rsv5,* confer resistance to the US SMV strains have been mapped to three chromosomes 2,13, and 14, respectively (Klepadlo *et al*., 2017; Saghai Maroof *et al*., 2008).

In China, a total of 22 SMV strains designated as SC1–SC22 were identified based on their responses to ten soybean differentials (Wang *et al.,* 2003, 2005; Guo *et al.,* 2005; Li *et al.,* 2010). Similar to the location of SMV resistance loci (*Rsv1*, *Rsv3*, *Rsv4* and *Rsv5*) identified in the US, the SMV resistance loci (mostly named *Rsc*) carried by Chinese soybean cultivars to SMV-SC strains have also been identified, and these loci are mostly located on chromosomes 2, 13, and 14 (Li *et al*., 2015; Ma *et al*., 2010, 2011; Zheng *et al*., 2014; Karthikeyan *et al*., 2018; Wang *et al*., 2018). Only one SMV resistance locus is different from the previous mapping results, *Rsc15*, a new locus located on chromosome 6 in cultivated soybean (cv.) RN-9 (Yang and Gai 2010; Ren *et al*., 2017).

Kefeng No.1 is a soybean cultivar that is resistant to multiple SMV strains. It’s reported that Kefeng No.1 carries many resistant loci, such as *Rsc5*, *Rsc7*, *Rsc8*, *Rsc10*, and *Rsc18A*, and these resistant loci are all located on chromosome 2 (Li *et al*., 2015; Ma *et al*., 2010, 2011; Zheng *et al*., 2014; Karthikeyan *et al*., 2018; Wang *et al*., 2018). Although these locaton regions in Kefeng No.1 overlap with each other, no resistant gene has been cloned and verified so far. Recently, Ishibashi *et al*. verified a SMV resistance gene, *Rsv4*, which is located on chromosome 2 in a broad-spectrum resistant cv. Peking (Ishibashi *et al*., 2019). *Rsv4* codes an RNase H family protein that mediates Peking’s resistance to multiple SMV strains by degrading double-stranded RNA (dsRNA). It is worth noting that the location region of *Rsv4* is adjacent to multiple SMV resistance loci of Kefeng No.1. Since both Kefeng No. 1 and Peking are both broad-spectrum SMV-resistant cultivars, and the resistant loci are located on chromosome 2, so will the resistance gene of Kefeng No. 1 mediate resistance to multiple SMV strains like *Rsv4*, or will there be many resistance genes in Kefeng No. 1 that mediate resistance to different strains? This issue has not yet been resolved.

In all virus-plant interaction systems involving typical or atypical R genes, a single viral protein corresponds to the R gene carried by the host. Replacing the viral protein changes the virulence of the virus on the host (Z Divéki *et al*., 2004; Hjulsager *et al*., 2006; Hull 2002; Padgett *et al*.,1997; Salvador *et al*., 2008). In previous research, after swapping SMV-G7 and SMV-G7d for P3 protein, it was found that the virulence of the two chimeric viruses on *Rsv1*-genotype soybean changed. Later, Hajimorad *et al*. found that SMV-N is avirulent on PI96983 (*Rsv1*-genotype soybean cultivar), and SMV-G7 is virulent on PI96983. Only when the HC-Pro and P3 of SMV-N are replaced with the corresponding proteins of SMV-G7 at the same time can the mutant gain the ability to infect PI96983 (Eggenberger *et al*., 2008; Hajimorad *et al*., 2006 and 2008; Khatabi *et al*., 2013; Wen *et al*., 2013). The results indicate that *Rsv1*-mediated resistance requires P3 and HC-Pro to induce, suggesting that *Rsv1*-genotype soybean may have different resistance genes (Wen *et al*., 2013). The virulence determinant causing *Rsv3*-mediated resistance has been mapped to CI. Replacement of CI of the virulence isolates with the counterparts from avirulence isolates caused ineffective infection of the recombinant virus on *Rsv3/Rsc4-3*-genotype soybeans (Seo *et al*., 2009; Zhang *et al*., 2009; Chowda-Reddy *et al*., 2011a; Yin *et al*., 2021). On *Rsv4*-genotype soybeans, only P3 was found to confer virulence of SMV via amino acid substitutions assay (Chowda-Reddy *et al*., 2011b; Khatabi *et al*., 2012; Ahangaran *et al*., 2013; Wang *et al*., 2015).

In this study, the F_2_-derived F_3_ (F_2:3_) and F_7:13_-derived recombinant inbred line (RIL) populations of soybean cv. Kefeng No.1 and cv. Nannong 1138-2 were used to locate the resistance gene on chromosome 2. VIGS assay showed that *Rsc3K* plays an important role in the resistance of Kefeng No.1 to SC3. Shuffling genes between virulence and avirulance SMV strains indicated that P3 of SMV is the virulence determinant of Kefeng No.1. Yeast two-hybrid (Y2H) and bimolecular fluorescent complementation (BiFC) experiments suggested that the P3 protein interacted with Rsc3K. In conclusion, we found that the resistance gene in the SMV resistant soybean cultivar Kefeng No.1 was consistent with the *Rsv4* gene in the SMV resistance soybean cultivar Peking previously reported, and both induced the resistance response by recognizing P3.

## Materials and methods

### Plant materials and mapping populations

Soybean cultivar Kefeng No.1 is resistant and cv. Nannong 1138-2 is susceptible to the SMV-SC3 strain. The 160 F_2:3_ and129 F_7:13_-derived RIL populations derived from a cross from Kefeng No.1 and Nannong 1138-2 were used to examine the inheritance of resistance and map the resistance gene *Rsc3K*. The mapping populations were planted in plastic pots (φ20×h20 cm) in an aphid-free net roomat 25°C . The soybean varieties and SMV isolates were provided by the National Center for Soybean Improvement (NCSI), Nanjing Agricultural University (NJAU), China.

### Inoculation and detection of SMV virus

SC3, an avirulent SMV strain to soybean cv. Kefeng No.1, and 1129, a virulent SMV isolate to Kefeng No.1. The inoculum was prepared by grinding infected fresh leaves of Nannong 1138-2 in 0.01 mol/L sodium phosphate buffer (approx. 3–5 mL per gram leaf tissue, pH 7.4) using mortars and pestles, and they were kept on ice until the inoculation was completed. The leaves for inoculation were rinsed with tap water after inoculation.

The unifoliate leaves of two populations were subjected to mechanical inoculation with SC3. At 7 dpi, soybean plants were subjected to mechanical inoculation secondly. Infection responses were observed weekly during the 40 dpi. Whereafter, the upper leaves with stable mosaic symptoms were used for virus detection by ELISA assay. And the segregation patterns of phenotypes in the mapping population were tested for the goodness-of-fit to Mendelian segregation ratio using the Chi-square test.

### DNA extractions, PCR process, and genetic analysis

The genomic DNA was extracted from the young leaves by using the NuClean Plant Genomic DNA Kit (Cwbio, Nanjing,China) and was stored at − 20°C. The primers of the SSR markers were synthesized by Genral Biological System (Nanjing, China) and used to detect polymorphisms between Kefeng No.1 and Nannong 1138-2.

Polymerase chain reactions (PCRs) were set in 10 μL volumes and the reaction mixtures included 1μL of genomic DNA (20 ng/μL), 5μL of 2 × Taq PlusMasterMix (Cwbio, Jiangsu,China), 0.5μL each of the forward and reverse primers (10 μM), and 3μL of double distilled water. The PCR conditions were as follows: preliminary denaturation at 94 °C for 2 minutes; 34 cycles of denaturation at 94 °C for 15 seconds (s) , annealing at 54 °C for 15 s, extension at 72 °C for 20 s; and cool at 4°C. All obtained PCR products were separated on 8 % polyacrylamide gels with silver staining (Wang *et al*., 2011b). The marker used to indicate the fragment length is 50bp DNA Ladder (TianGen, BeiJing, China) or DL5000 (Takara, Dalian, China).

The genetic distances between markers were calculated by Join Map 4.0 (Van Ooijen *et al*., 2006). The distances between markers and the resistance gene were calculated with Kosambi genetic distances (Kosambi 2011). Map Chart v. 2.1 (Voorrips 2002) was used to draw the linkage map.

### Construction of segregating pools and fine-mapping the *Rsc3K* gene

BSA was performed for mapping the *Rsc3K* gene. The leaves of R-pool and S-pool individuals of the RIL population were collected and extracted DNA. The R/S-pool was made by mixing equal amounts of DNA from 20 RIL individuals, respectively.

Forty SSR markers reportedly linked to several SMV-resistance loci were to determine whether they were polymorphic in SMV-SC3-resistant parent and susceptible parent, from which 6 polymorphic SSR markers (BARCSOYSSR_02_0478, BARCSOYSSR_02_0555, BARCSOYSSR_02_0610, BARCSOYSSR_02_0620, BARCSOYSSR_02_0632 and BARCSOYSSR_02_0670) were employed to construct a genetic map. Four additional SSR markers (BARCSOYSSR_02_0614, BARCSOYSSR_02_0615, BARCSOYSSR_02_0617, BARCSOYSSR _02_0619) were selected for genotype analysis of recombined lines. All primers for SSR markers werelisted in Supplemental Table 1.

### Candidate genes prediction and construction of silencing vectors

Putative candidate gene models in the identified flanking region were predicted according to the NCBI (https://www.ncbi.nlm.nih.gov/; Glycine max Wm82. a2.v1; verified December, 2016). Primers for the amplification of candidate genes were designed from the respective genome sequences using Primer Premier 5.0 software (Premier Biosoft, Palo Alto, USA). All primer sequences were listed in Supplemental Table 1. Genes that were successfully amplified were sequenced (Generalbio, Anhui, China). Sequence alignment and analysis were performed by using BioXM 2.7 software (http://www.biosoft.net)(NJAU, China)(Huang and Zhang 2004).

The potential resistance candidate genes were verified by VIGS. The method of constructing BPMV silencing vectors and plasmids inoculation followed Zhang (Zhang *et al*., 2010). The pBPMV-IA-V2-R2 was cut by BamHI and SalI (NEB, Beijing, China) and the fragments of candidate genes were amplified from Kefeng No.1 plants. Then the homologous recombination method was used to connect with liner vector and fragments.The constructed vectors were mixed with pBPMV-IA-R1M plasmids respectively, and the mixture inoculated on Nannong 1138-2 plants by mechanical inoculation.

### Detect the silencing efficiency of candidate genes and the accumulation of SMV in silencing plants

When the Nannong 1138-2 plants inoculated with the silencing vectors showed mosaic symptom, the diseased leaves were used to extract mRNA. Total RNA was extracted using TRIzol reagent (Invitrogen, Carlsbad, USA), and cDNA was synthesized using HiScript® II QRT SuperMix (VazymeBiotech, Nanjing, China). Then RT-PCR was performed to detect the BPMV vectors containing the inserted fragments. Sequences of detection primers of BPMV-R2-C2F/R were listed in Supplemental Table 1. The diseased plants with different silencing vectors were inoculated onthe unifoliate leaves of Kefeng No.1, and SMV-SC3 was inoculated on the first trifoliate leaves of Kefeng No.1. Two weeks after inoculation with SC3, the diseased leaves of Kefeng No.1 plants were collected to detect the silencing efficiency of the corresponding silencing vectors and the SMV accumulation. The relevant detection of Kefeng No.1 plants silenced-*Gm02R* inoculated with other SMV strains (SC5, SC7, SC8, SC10, and SC18) were consistent with the above steps. The ELISA assay was used to verify the susceptibility of the samples and the qRT-PCR was used to analyze the silencing efficiency of candidate genes and the accumulation of SMV. Relative quantification of gene expression was carried out with the 2^-ΔΔ^Ct method. Each treatment contained three samples and was repeated three times.

### Sequencing and alignment SMV SC3 and 1129

When the unifoliate leaves of Nannong 1138-2 had fully unfolded, they were inoculated with SMV-SC3 and SMV-1129, respectively. At 7dpi, the diseased leaves of Nannong 1138-2 were used to extract total RNA. Based on RT-PCR, the nucleotide sequences of SMV-SC3 and SMV-1129 were obtained. All primers used to amplify the genome sequences of the two isolates were listed in Supplementary Table 1.

The nucleotide and amino acid sequences of SMV-SC3 (GenBank, MH919384) and SMV-1129 were aligned by Snapgene 4.1.9 (GSL Biotech LLC, Chicago, USA).

### Construction of recombinant and site-directed clones

The method of constructing SMV infectious clones of SMV-SC3 and SMV-1129 was followed Yin *et al*. 2019. In short, the nucleotide sequences of SMV-SC3 and SMV-1129 were amplified into five segments and all the fragments and the linearized vector pCB301 (StuI and SmaI)(NEB, Beijing, China) were mixed to transform the yeast strain W303-113.

The process of constructing recombinant clone is the same as the above method. Taking pSMV-SC3-R1 as an example, first, five fragments of SMV-SC3 and SMV-1129 were amplified, respectively. Then the recombinant fragment between 6K2 in SC3 and NIa-Vpg in SMV-1129 was generated by overlapping PCR. Finally, all fragments and vectors were mixed into yeast strain W303-113. Two combinant fragments between 6K1 in SMV-SC3 and CI in SMV-1129, HC-Pro in SMV-SC3 and P3 in SMV-1129 were also generated by overlapping PCR and to construct two other recombinant clones pSMV-SC3-R2 and pSMV-SC3-R3.

To identify mutations essential for breaking Kefeng No.1 resistance, site-directed mutagenesis was performed. Two site-directed mutations (G1055R and E1126V) were introduced into pSMV-SC3 to produce clones pSMV-SC3_G1055R_ (G1055R) and pSMV-SC3_E1126V_ (E1126V). Furthermore, a revertant mutant clone, pSMV-1129_R1055G_ (R1055G), was constructed, whose amino acid sequences were consistent with SMV-1129 except for 1055aa.

We screened the recombinant clones by PCR and sequencing methods. All primer sequences used to construct the recombinant vectors were listed in Supplementary Table 1. The method of inoculation and detection of the recombinant cloned plasmids were the same as that of the VIGS vectors.

### The Y2H assay between soyeban proteins and SMV P3

*Rsc3K* from Kefeng No.1 and SMV *P3* from both SMV-SC3 and SMV-1129 were cloned into the prey vector pPR3 (pPR3-Rsc3K) and the bait vectors pBT3 (pBT3-SC3 P3 and pBT3-1129 P3), respectively. The resultant vectors were co-transformed into yeast strain NMY51 and cultured overnight at 30 °C for 4 days on SD/–Leu/–Trp (double dropout) and SD/-Trp/-Leu/-His/-Ade (quadruple dropout) medium. The density of the yeast colonies was adjusted such that the OD_600_ was 0.2, and they were then transferred to the same solid medium and diluted 1/10, 1/100 and1/1000 to produce a concentration gradient that was used to determine the mutual affinity between Rsc3K and the two P3 proteins. The two genes, *LOC100526921* and *LOC100812666* from cv. Williams 82 were constructed into the prey vector pPR3 (pPR3-LOC100526921 and pPR3-LOC100812666), and then transferred into yeast cells with the two vectors pBT3-SC3 P3 and pBT3-1129 P3, respectively. The transformation method was the same as above. The operation steps should be followed the protocol of DUAL membrance Kit 3 (Dualsystems Biotech, Switzerland).

### The BiFC assay between soybean proteins and SMV P3

The BiFC assay was used to confirm the interaction between Rsc3K from Kefeng No.1, LOC1000526921 and LOC100812666 from Williams 82 and P3 of two SMV isolates in *Agrobacterium*-infiltrated *N. benthamiana* plants. Among them, three soybean genes were cloned into the pEarleyGate202-YN vector (Rsc3K-NYFP), and the two *P3* genes were cloned into the pEarleyGate201-YC vectors (SC3 P3-CYFP and 1129 P3-CYFP). All constructs were transformed into *Agrobacterium tumefaciens* strain EHA105, and the OD_600_ was adjusted to 1 with infiltration buffer. The infiltration buffers of Rsc3K (or LOC1000526921 and LOC100812666 )-NYFP , SC3 P3-CYFP or 1129 P3-CYFP and PM marker were mixed in equal amounts and agroinfiltrated into 4-week-old *N. benthamiana* leaves. At 3dpi, the agroinfiltrated leaves were observed by laser scanning confocal microscopy (Zeiss LSM 780, Germany).

### Protein model prediction and Phylogenetic tree analysis

The predicted protein model of Rsc3K, LOC1000526921, LOC100812666, SMV-SC3 P3, and the mutatant P3 (Q1034K and G1055R) were obtained through TMHMM2.0 (https://services.healthtech.dtu.dk/service.php?TMHMM-2.0 )(Krogh *et al*., 2001). The transmembrane structure protein model of SMV-SC3 P3 and the mutatant P3 (Q1034K and G1055R) were confirmed by HMMTOP (http://www.enzim.hu/hmmtop/html/submit.html)(Tusnády and Simon, 1998 and 2001).

Phylogenetic tree analysis of Chinese SMV isolates were conducted in MEGA7 (Kumar *et al*., 2016), and the phylogenetic tree was constructed using Neighbor-Joining method.

## RESULTS

### *Rsc3K* is the dominant resistance gene of soybean cv. Kefeng No.1 to SC3

To determine the inheritance pattern of SMV resistance in soybean cv. Kefeng No.1, F_2:3_ and F_7:13_-derived RIL populations obtained from the cross of resistant Kefeng No.1 and susceptible Nannong 1138-2 were inoculated with SMV-SC3. Forty days after inoculation with SMV-SC3, the F_2:3_ population showed a divergence of resistance, including 25 resistant lines, 46 susceptible lines, and 89 heterozygous lines (Table 1). The RIL population contained 64 resistant lines and 65 susceptible lines (Table 1). The F_2:3_ and RIL populations segregated in a 1R: 2 Segregate (Seg) :1S and 1R:1S ratio, respectively. As a control, all 20 Kefeng No. 1 plants showed resistance, while 20 Nannong 1138-2 plants showed susceptibility (Table 1). These results suggest that a dominant gene, designated as *Rsc3K* controls resistance of Kefeng No.1 to SC3.

**Table 1.**
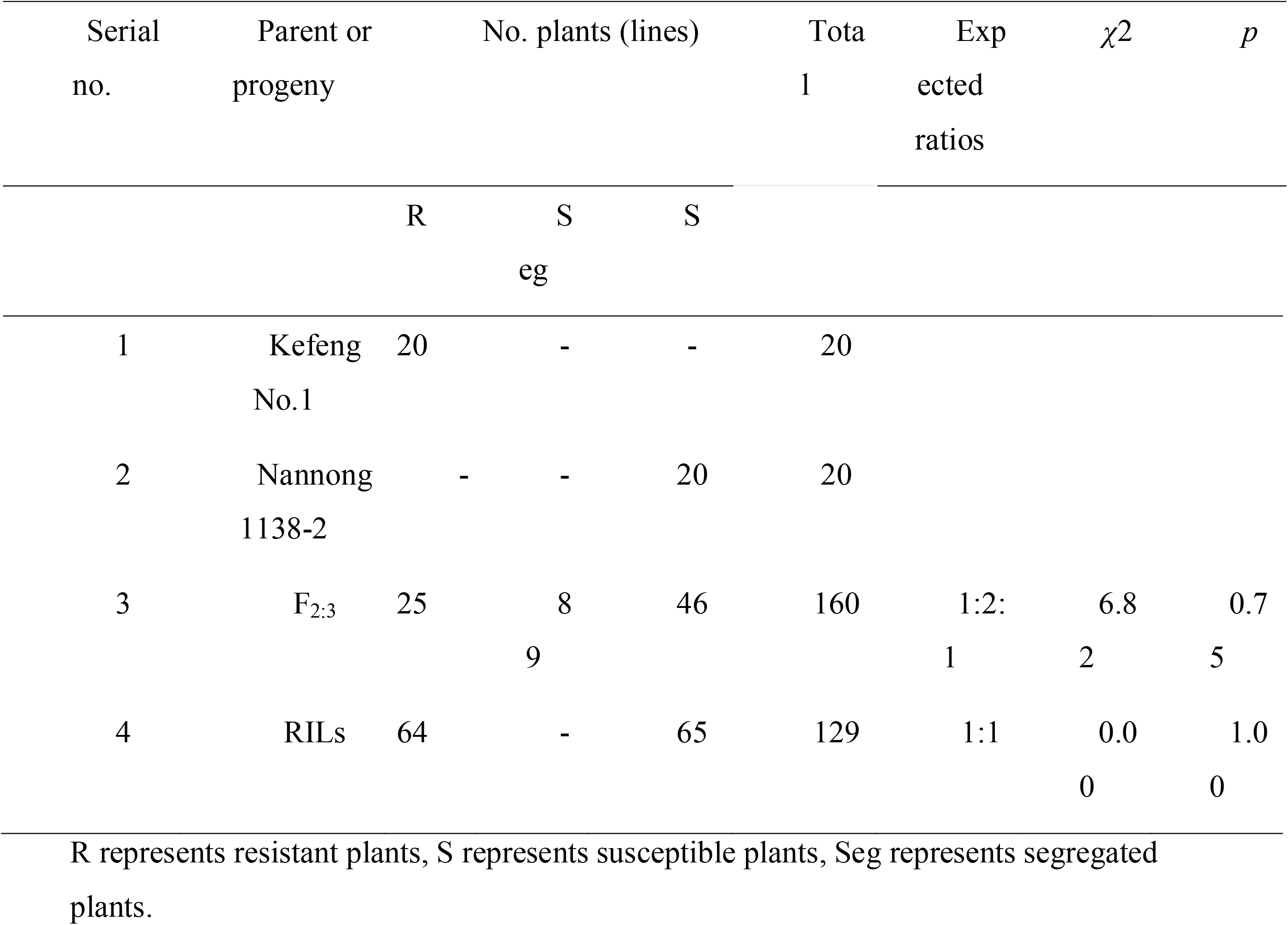
Genetic of resistance to SMV-SC3 in Kefeng No.1.

### *Rsc3K* was fine-mapped on chromosome 2 of Kefeng No.1

In previous reports, resistance loci to SMV in soybean were located on chromosomes 2, 6, 13, and 14. From 40 Simple Sequence Repeat (SSR) markers linked to those loci, we screened out 9 markers with polymorphism between the resistant parent Kefeng No. 1 and the susceptible parent Nannong 1138-2. Then we used these 9 markers to locate *Rsc3K* by Bulked Segregant Analysis (BSA) method. Based on genotype analysis, six SSR markers were identified to be linked to *Rsc3K*, namely BARCSOYSSR_02_0478, BARCSOYSSR_02_0555, BARCSOYSSR_02_0610, BARCSOYSSR_02_0620, BARCSOYSSR_02_0632, and BARCSOYSSR_02_0670. These SSR markers are located on chromosome 2, suggesting that the resistance gene is located on chromosome 2. Based on the genotypic information of 129 individuals of the RIL population obtained by using these markers, we constructed a linkage map. The total length of the linkage map was 45.3 centimorgan (cM) and the resistance gene was mapped to an interval between BARCSOYSSR_02_0610 and BARCSOYSSR_02_0620 (Figure 1). The genetic distances for the resistance gene were 4.3 and 3.0 cM to the adjacent markers. In order to narrow down the mapping interval, four additional SSR markers, BARCSOYSSR_02_0614, BARCSOYSSR_02_0615, BARCSOYSSR_02_0617, and BARCSOYSSR_02_0619 were used to identified recombination lines between markers BARCSOYSSR_02_0610 and BARCSOYSSR_02_0620. Amongthem, 10 recombination lines (named K1-K10) were filtrated, and one line K10 was found to have the recombinted site between markers BARCSOYSSR_02_0614 and BARCSOYSSR_02_0615 (Table 2). We observed that K10 was susceptible to SMV-SC3. Since K10 had the same genotypes as resistant parent at markers BARCSOYSSR_02_0610 and BARCSOYSSR_02_0614 , and the same genotypes as susceptible parent at markers at other four markers (Table 2), we conclude that *Rsc3k* is located at the downstream of marker BARCSOYSSR_02_0614 (Table 2). Thus, the candidate region was delimited to a physical interval of approximately 90 kb flanked by BARCSOYSSR_02_0614 and BARCSOYSSR_02_0620.

**Figure 1.**
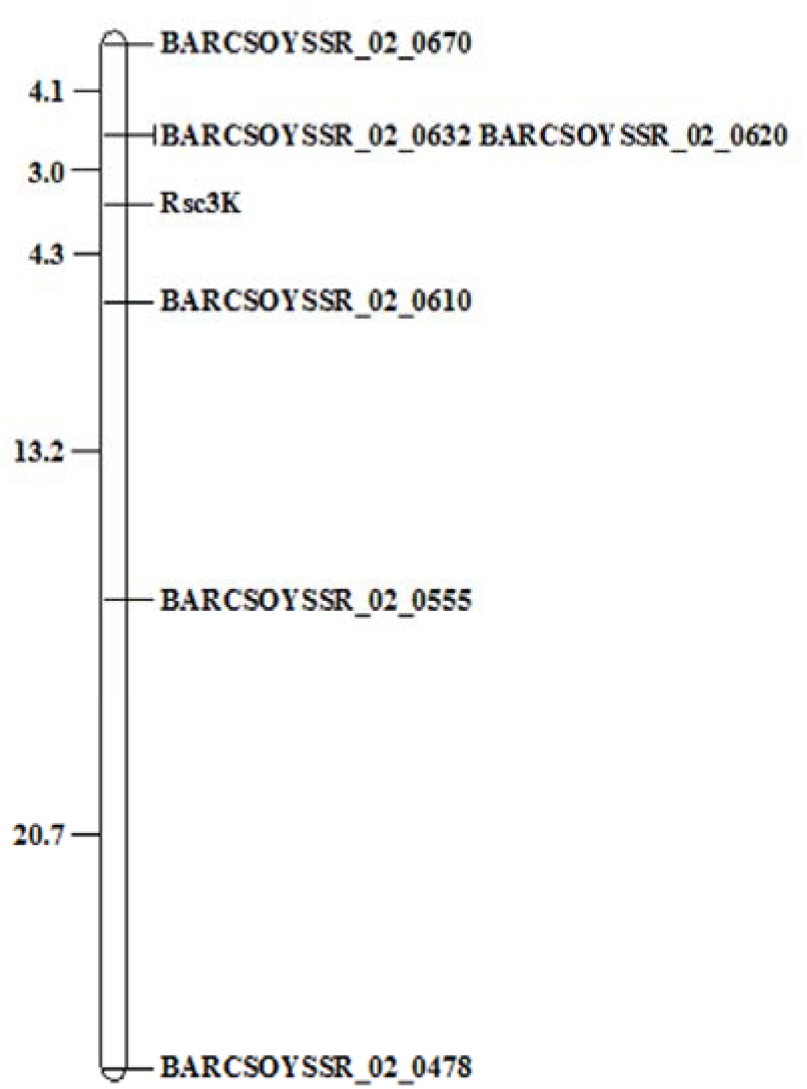
*Rsc3K* was located on chromosome 2 in soybean cultivar Kefeng No.1. By using the F_7:13_-derived RIL population derived from a cross between soybean cv. Kefeng No. 1 (resistant) and cv. Nannong 1138-2 (susceptible), we mapped the resistance gene (*Rsc3K*) of Kefeng No. 1 to SMV-SC3 on chromosome 2. The genetic linkage map of the *Rsc3K* gene was plotted, map distances (cM) between neighboring markers are on the left with marker names on the right.

**Table 2.**
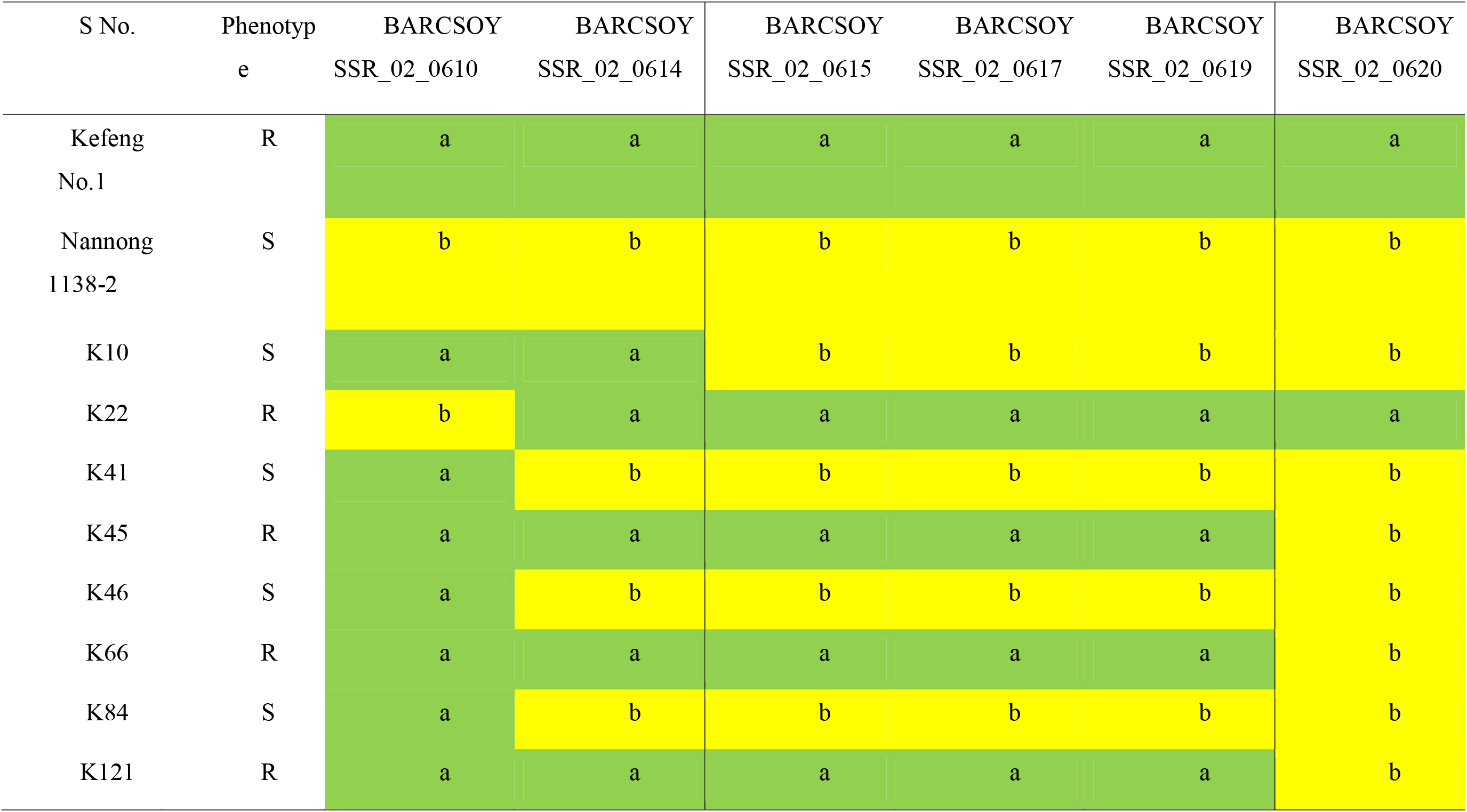
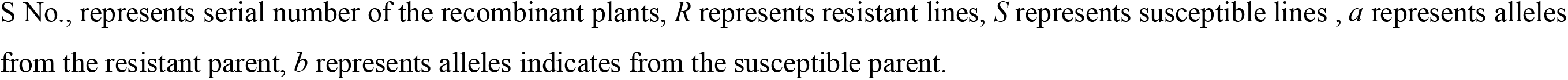
Phenotypes and genotypes of recombinant lines showing the recombinant breaking points.

| S No. | Phenotype | BARCSOY<br>SSR_02_0610 | BARCSOY<br>SSR_02_0614 | BARCSOY<br>SSR_02_0615 | BARCSOY<br>SSR_02_0617 | BARCSOY<br>SSR_02_0619 | BARCSOY<br>SSR_02_0620 |
| --- | --- | --- | --- | --- | --- | --- | --- |
| Kefeng<br>No.1 | R | a | a | a | a | a | a |
| Nannong<br>1138-2 | S | b | b | b | b | b | b |
| K10 | S | a | a | b | b | b | b |
| K22 | R | b | a | a | a | a | a |
| K41 | S | a | b | b | b | b | b |
| K45 | R | a | a | a | a | a | b |
| K46 | S | a | b | b | b | b | b |
| K66 | R | a | a | a | a | a | b |
| K84 | S | a | b | b | b | b | b |
| K121 | R | a | a | a | a | a | b |

S No., represents serial number of the recombinant plants, *R* represents resistant lines, *S* represents susceptible lines, *a* represents alleles from the resistant parent, *b* represents alleles indicates from the susceptible parent.

### Four candidate genes of the localization segment were screened out

There are 10 putative genes in the 90 kb mapping interval according to Williams 82 genomic database. Among them, one gene has no clear functional annotation, one belongs to non-coding RNA (ncRNA), and the others are involved in disease defense, transcription, and metabolism (Table 3). Based on the predicted gene models, two genes (*LOC100811049* and *LOC100811588* ) are MADS transcription factors involved in biotic and abiotic stresses. Two genes (*LOC106797235* and *LOC100527708*) are related to ubiquitination and hydrolase metabolic processes, respectively. The other four genes, may play roles in disease defense, two of them (*LOC100776324* and *LOC100776859*) are receptor kinase, one (*LOC100778452*) is a molecular chaperone, and one (*LOC100812666*) has mediates disease resistance by degrading double-stranded RNA. Finally, these eight genes were selected as candidate genes and cloned from Kefeng No.1(resistant cultivar) plants for comparison with alleles of Williams 82 (susceptible cultivar).

**Table 3.** Annotation of 10 cadidate genes in the *Rsc3K* mapping interval.

| Ser<br>ial no. | Genes | Annotation | Functional<br>classification |
| --- | --- | --- | --- |
| 1 | *LOC100811049* | Agamous-like<br>MADS-box protein MADS2 | Transcription<br>factor |
| 2 | *LOC100811588* | MADS-box<br>transcription factor | Transcription<br>factor |
| 3 | *LOC106797235* | Probable E3<br>ubiquitin-protein ligase<br>RHB1A | Metabolism |
| 4 | *LOC100527708* | Adenine nucleotide<br>alpha hydrolases-like<br>superfamily protein | Metabolism |
| 5 | *LOC112999319* | Non-coding RNA |  |
| 6 | *LOC100776324* | Leucine-rich repeat<br>receptor-like protein kinase | Disease<br>defense |
| 7 | *LOC100776859* | Receptor-like protein<br>kinase FERONIA | Disease<br>defense |
| 8 | *LOC100812666* | Putative ribonuclease H<br>protein At1g65750-like | Disease<br>defense |
| 9 | *LOC100526921* | Uncharacterized |  |
| 10 | *LOC100778452* | OmpH and DnaJ<br>domain-containing protein | Disease<br>defense |

The candidate genes in the *Rsc3K* location region were listed, according to the Williams 82 soybean reference genome annotation Glycine\_max\_v2.1 (accessible at https://www.ncbi.nlm.nih.gov/) (verified 24th July 2018).

By amplifying the DNA from Kefeng No.1, seven candidate genes correspond to the genome of Williams 82. Based on sequence comparison, DNA sequences of *LOC100811049* and *LOC100778452* are identical within alleles from Kefeng No.1 and Williams 82. *LOC106797235* and *LOC100527708* encode proteins that contain same amino acids, despite differences on the DNA level. *LOC100811588*, *LOC100776324*, and *LOC100776859* in the Kefeng No.1 genome contains nonsynonymous SNP mutations or frameshift mutations comparing to their alleles in William 82 (Table 4).

**Table 4.**
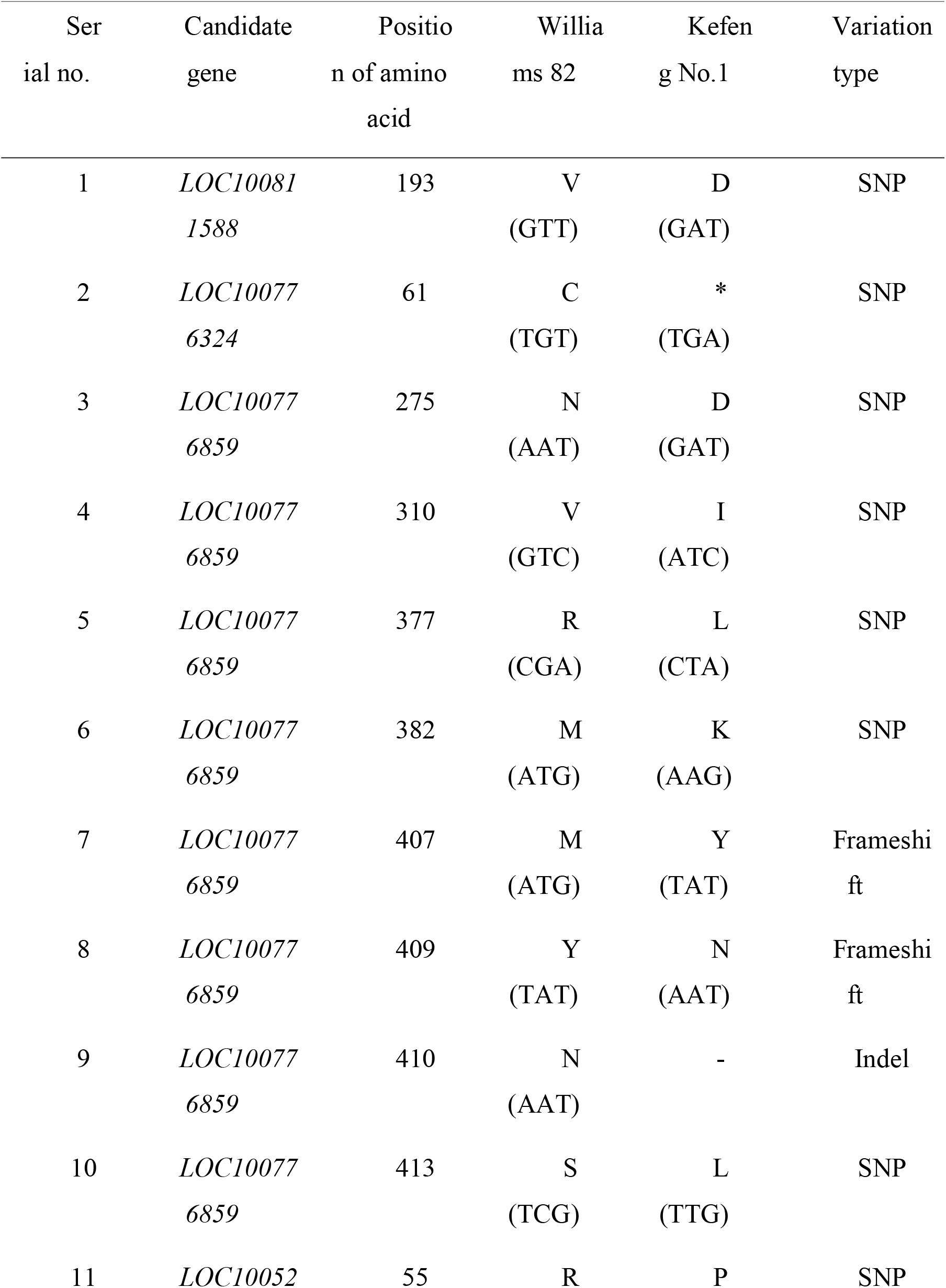
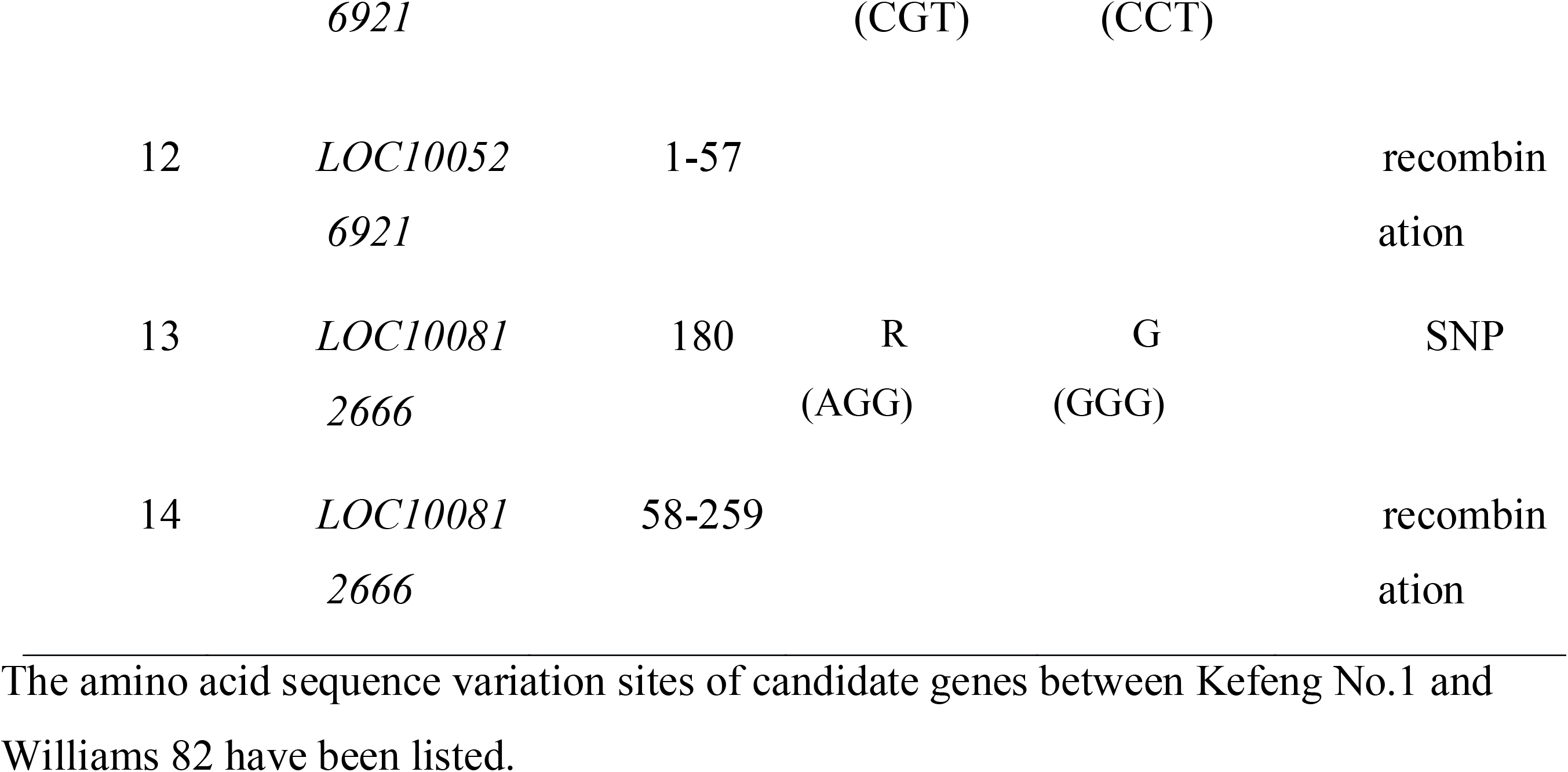
Sequence variation of Rsc3K candidate genes in Kefeng No.1.

To our suprise, the coding sequence of *LOC100812666* could not be amplified from Kefeng No.1 genomic DNA by using the primers designed according to the Williams 82 genome. We speculated that there might be a large variation of *LOC100812666* allele in Kefeng No.1. So we designed primers at 1kb, 2kb, 3kb, and 4kb upstream or downstream of the *LOC100812666*, according to the Williams 82 reference genome. A fragment of approximately 1kb was amplified from Kefeng No.1 by using the forward primer at 4kb upstream of *LOC100812666* and reverse primer that recognize the 3’-end of *LOC100812666* gene (Figure 2A). Meanwhile, all PCR fragments of expected size could be amplified from Williams 82 (Figure 2A and 2B). By analyzing the sequence of the amplified DNA fragments, we found that an internal deletion happens in the Kefeng No.1, comparing to the William 82 reference genome. This deletion event results a recombinant gene product that has N-terminal 57 amino acids originated from reference gene *LOC100526921* and C-terminal 202 amino acids from reference gene *LOC100812666* (Table 4, Figure 3). We temporarily named this recombinant gene *Gm02R*.

**Figure 2.**
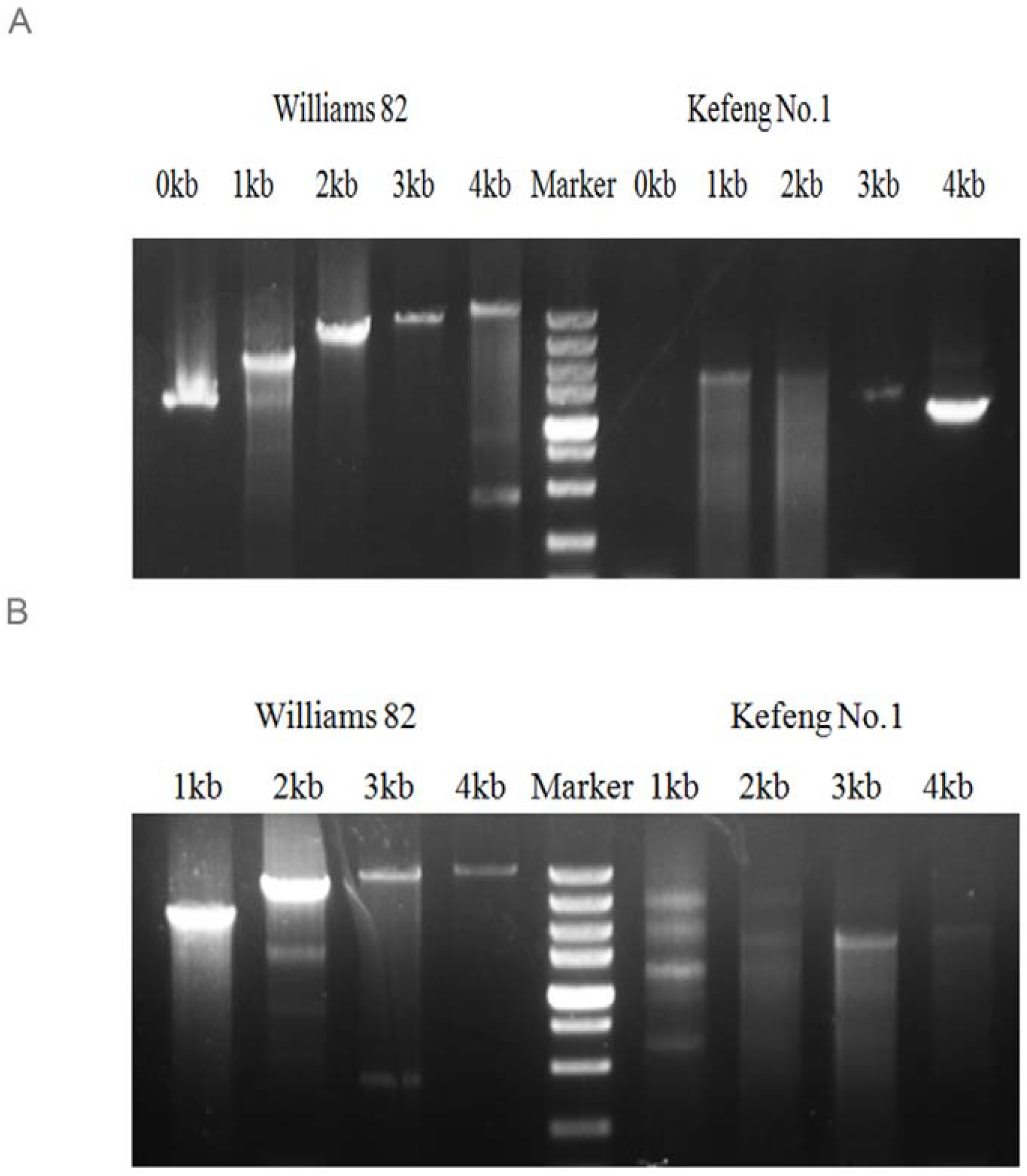
There is a large deletion in the mapping region of *Rsc3K* in Kefeng No. 1. By amplifying the candidate genes located in the mapping region, we found that there is a deletion in the genome of Kefeng No. 1, which is located upstream of *LOC100812666*. Images of amplified fragments upstream and downstream of *LOC100812666* from William 82 and Kefeng No. 1 were shown. Marker is DL5000, where the brightest band is 1000bp and the largest band is 5000bp. (A) The fragments of 0kb, 1kb, 2kb, 3kb, and 4kb upstream of *LOC100812666* were amplified in the genomes of William 82 and Kefeng No. 1. (B) The fragments of 1kb, 2kb, 3kb, and 4kb downstream of *LOC100812666* were amplified in the genomes of William 82 and Kefeng No. 1.

**Figure 3.**
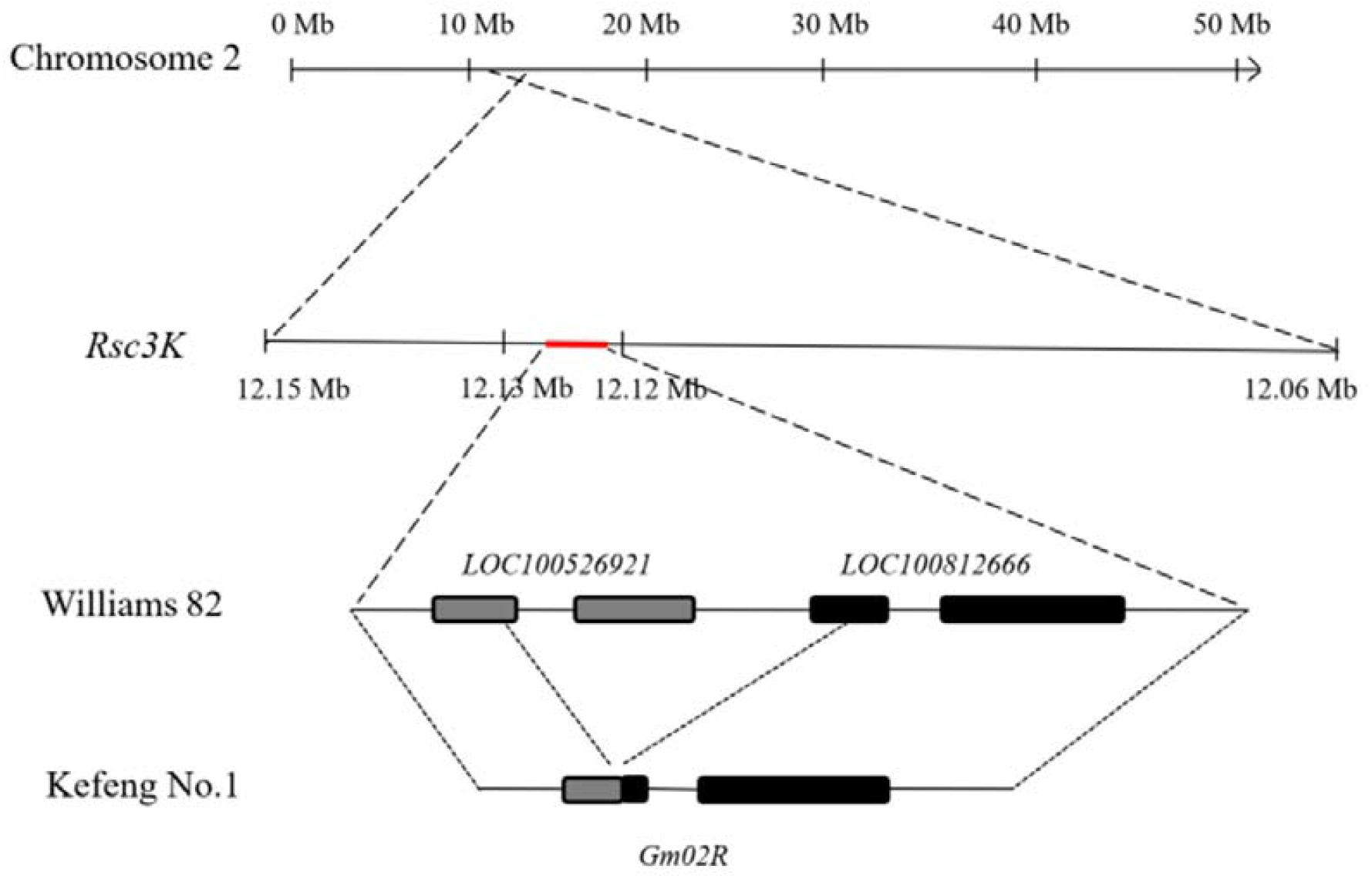
*Gm02R* in Kefeng No.1 is recombined from *LOC100526921* and LOC100812666. The gene structures of Gm02R in Kefeng No.1 and LOC100526921 or LOC100812666 in Williams 82 are shown. Gray bars represent the exons of *LOC100526921*. Black bars represent the exons of *LOC100812666*. The recombinant gene *Gm02R* in Kefeng No.1 is drew by gray and black bars. Black lines are untranslated regions.

### *Gm02R*-silenced plants showed more sensitive symptoms after inoculation with SMV strains

The three genes (*LOC100811588*, *LOC100776324*, and *LOC100776859*) that differs between Kefeng No.1 and William 82 and *Gm02R* were selected for the BPMV based VIGS assay to identify their role in SMV resistance (Zhang *et al*., 2010). The silencing vectors constructed were named as Si0201(silenced for *LOC100811588*), Si0202 (silenced for *LOC100776324*), Si0203 (silenced for *LOC100776859*), and Si0204 (silenced for *Gm02R*). The diseased plants with different silencing vectors were inoculated on the unifoliate leaves of Kefeng No.1, and SC3 was inoculated on the first trifoliate leaves of Kefeng No.1. Two weeks after inoculation of SMV-SC3, Kefeng No.1 plants inoculated with different silencing vectors previously showed different degrees of mosaic and shrinkage, and the *Gm02R*-silenced plants showed more severe mosaic and shrinkage than the control plants (inoculated with empty vector, pBPMV-V2 )( Figure 4A). The other genes-silenced plants, were similar mosaic symptoms to pBPMV-V2 plants (Figure 4A). The accumulation of SMV in infected leaves was detected by double antibody sandwich enzyme-linked immunosorbent assay (DAS-ELISA). The result of ELISA assay showed that the SMV accumulation in plants silenced for *Gm02R* was significantly higher than pBPMV-V2 plants (Figure 4B). However, the SMV accumulation in other plants silenced for *LOC100811588*, *LOC100776324*, and *LOC100776859* were similar to that of the pBPMV-V2 plants (Figure 4B). The transcript level of *Gm02R* was detected by qRT-PCR. The mRNA level of *Gm02 R* reduced by 77.2% in *Gm02R*-silenced plants compared to pBPMV-V2 plants (Figure 4C). This showed that *Gm02R* was effectively silenced in Kefeng No.1. Therefore, we speculated that *Gm02R* is an important gene in the SMV resistance process mediated by *Rsc3K*.

**Figure 4.**
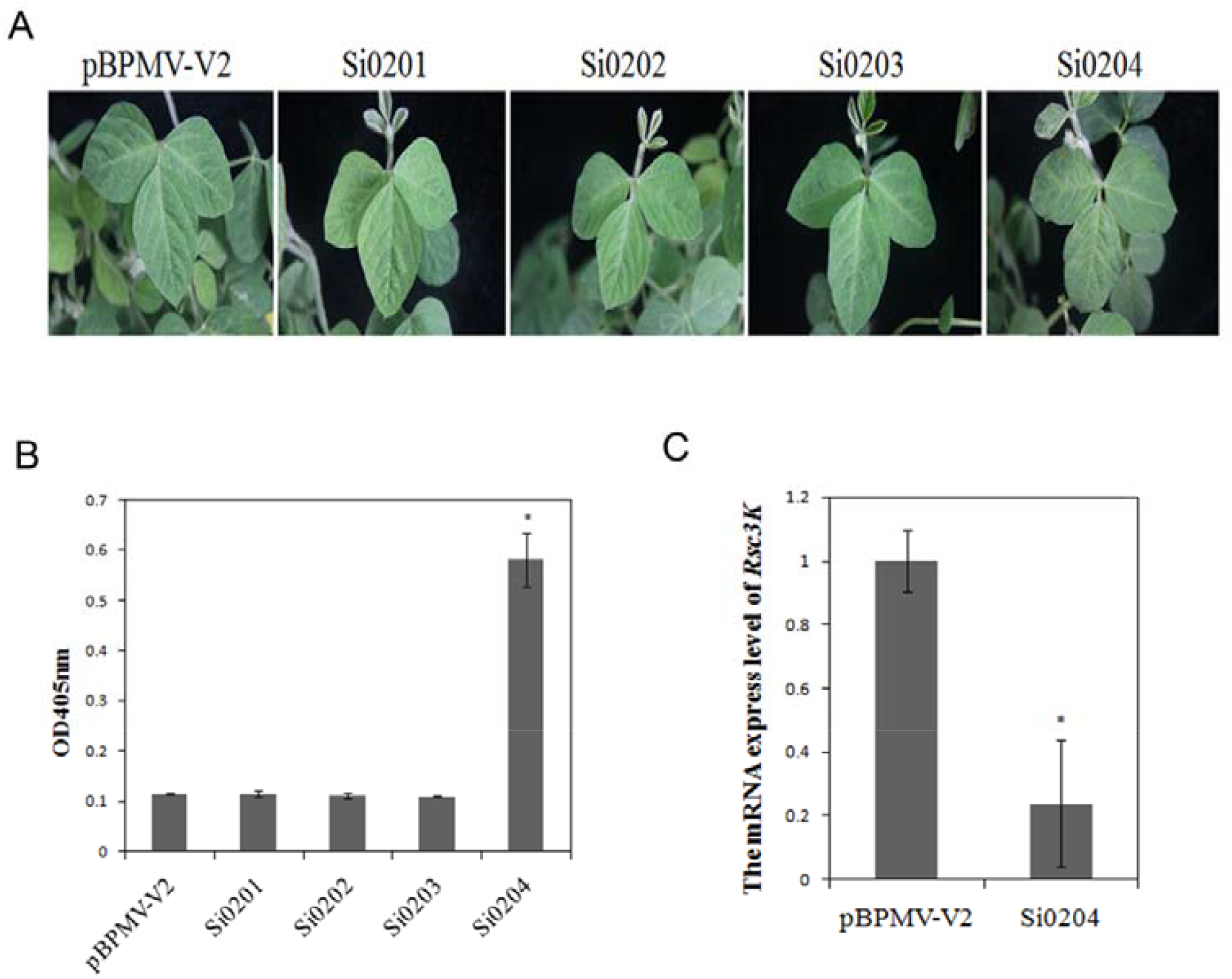
*Gm02R*-silenced plants were susceptive to SMV-SC3. All Kefeng No.1 plants were inoculated with BPMV-VIGS vectors on unifoliate leaves. At 7dpi, we inoculated SMV-SC3 on trifoliate leaves of genes-silenced plants. For convenience, we named the plants of silence candidates (*LOC100811588*, *LOC100776324*, *LOC100776859*, and *Gm02R*) as Si0201, Si0202, Si0202, and Si0204. In this experiment, plants inoculated with pBPMV-V2 were used as controls. Phenotypic observation and detection assay were performed at 14 dpi with SMV-SC3 inoculation. (A) The phenotypic responses on pBPMV-V2, Si0201, Si0202, Si0202, and Si0204 plants. (B) The SMV accumulation of pBPMV-V2, Si0201, Si0202, Si0202, and Si0204 plants were detected by ELISA assay. Each experiment has three replicate plants, with the standard errors indicated. Asterisks denote significant differences in the accumulation of SMV compared to that in control plants, t-test, p < 0.001. (C) The mRNA expression level of *Gm02R* in pBPMV-V2 and Si0204 plants were tested. Each experiment has three replicate plants, with the standard errors indicated. Asterisks denote significant differences in the expression of *Gm02R* mRNA compared to that in pBPMV-V2 plants, t-test, p < 0.001.

Kefeng No.1, a broad-spectrum resistant cultivar to SMV, showed symptomless after inoculation with SMV strains SC5, SC7, SC8, SC10, and SC18 (Figure 5A). So after silencing *Gm02R*, would Kefeng No.1 plants be susceptible to these strains? For this purpose, we inoculated *Gm02R*-silenced plants with SMV strains SC5, SC7, SC8, SC10, and SC18, and collected disease leaves for ELISA detection at 14 days post-inoculation (dpi). The results showed that these SMV strains could all infect Kefeng No.1 plants silenced for *Gm02R* (Figure 5B). In contrast, Kefeng No.1 plants can not be infected by SMV strains SC5, SC7, SC8, SC10, and SC18 (Figure 5B). These indicated that silencing of *Gm02R* in Kefeng No.1 contributed to SMV infection in Kefeng No.1. The VIGS assay demonstrated that *Gm02R* plays a key role in the resistance of Kefeng No.1 to SMV-SC3 and many other SMV strains. Therefore, we conclude that *Gm02R* might be responsible for *Rsc3K*, and *Rsc3K* may mediate the broad-spectrum resistance of soybean cv. Kefeng No.1 to SMV.

**Figure 5.**
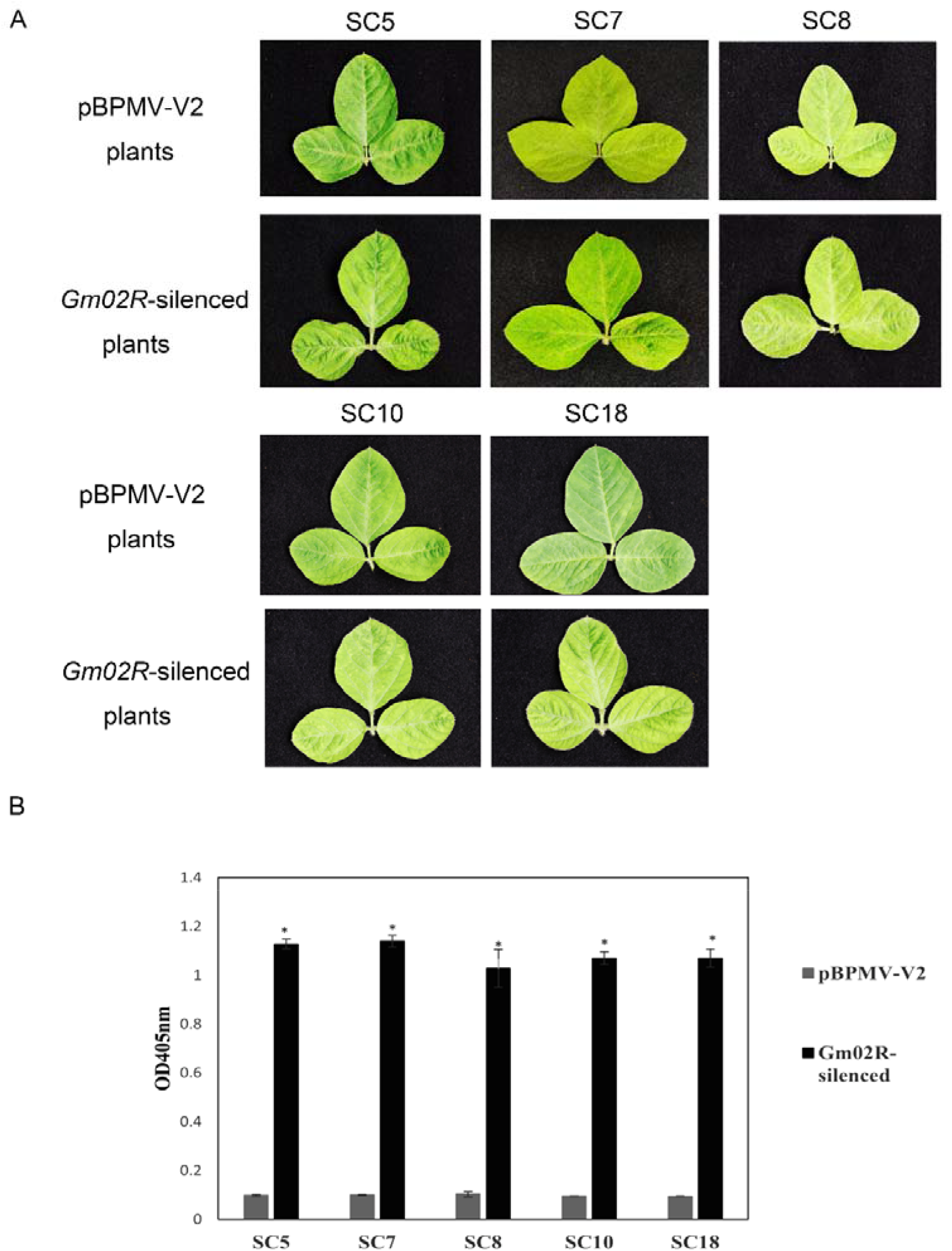
*Gm02R*-silenced plants were susceptive to other SMV strains (SMV-SC5, SMV-SC7, SMV-SC8, SMV-SC10, and SMV-SC18). The *Gm02R*-silenced and pBPMV-V2 plants were inoculated with SMV-SC5, SMV-SC7, SMV-SC8, SMV-SC10, and SMV-SC18. Phenotypic observation and detection assay were performed at 14 dpi with SMV strains inoculation. (A) The phenotypic responses on pBPMV-V2 and *Gm02R*-silenced plants with SMV-SC5, SMV-SC7, SMV-SC8, SMV-SC10, and SMV-SC18 inoculation, respectively. (B) The SMV accumulation of pBPMV-V2 and *Gm02R*-silenced plants were detected by ELISA assay. Each experiment has three replicate plants, with the standard errors indicated. Asterisks denote significant differences in the accumulation of SMV compared to that in control plants, t-test, p < 0.001.

### P3 is the virulence determinant of SMV on Kefeng No.1

We screened several SMV isolates and found that isolate 1129 can cause typical symptoms on soybean cv. Kefeng No.1, including prominent leaf mosaic and shrinkage (Figure 7A and 7C).

The whole genomes of SMV SC3 and 1129 were cloned and sequenced. Both viruses had a genome of 9588 nucleotides, and they were 99 % identical at both nucleotide and amino acid sequence levels. Each genome contains an ORF of 9201 nucleotides-long, which encodes a polyprotein with 3067 amino acids. There were 18 nucleotides and 8 amino acid differences between SC3 and 1129. The amino acid differences were located in six viral proteins, HC-Pro, P3, 6k1, CI, NIa-Pro, and CP (Table 5, Figure 6A). Therefore, we constructed recombinant clones using the avirulent SC3 and the virulent 1129 to find virulence determinants of SMV on Kefeng No.1. Three recombinant clones were generated, namely pSMV-SC3-R1, pSMV-SC3-R2, and pSMV-SC3-R3 (Figure 6A).

**Figure 6.**
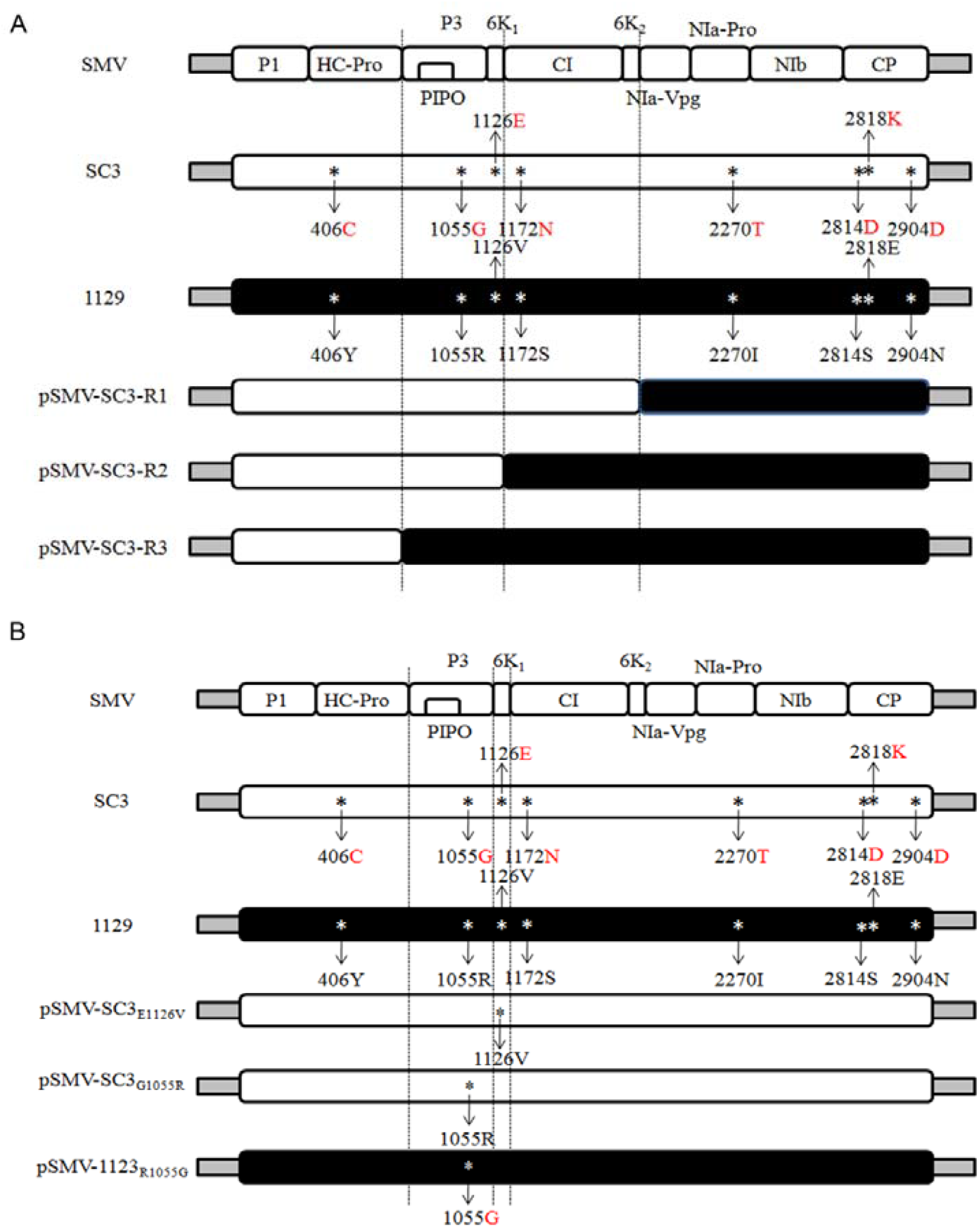
Recombinant and point mutagenesis clones were constructed by using the virulent SMV-1129 and avirulent SMV-SC3 on Kefeng No.1. The 11 mature proteins of SMV are marked in the figure, and the amino acids that differ between SMV-SC3 and SMV-1129 are represented by red and black letters, respectively. Gray bars represent the sequence of the vector, white bars represent the sequence of the recombinant clones consistent with the SC3, and black bars represent the sequence of the recombinant clones consistent with the 1129. Asterisks represent the mutation sites. (A) Schematic representation of the genomic maps of three recombinant clones (pSMV-SC3-R1, pSMV-SC3-R2, and pSMV-SC3-R3). (B) Schematic representation of the genomic maps of point mutagenesis clones (pSMV-SC3_G1055R_, pSMV-SC3_E11126V_, and pSMV-1129_R1055G_).

**Table 5.**
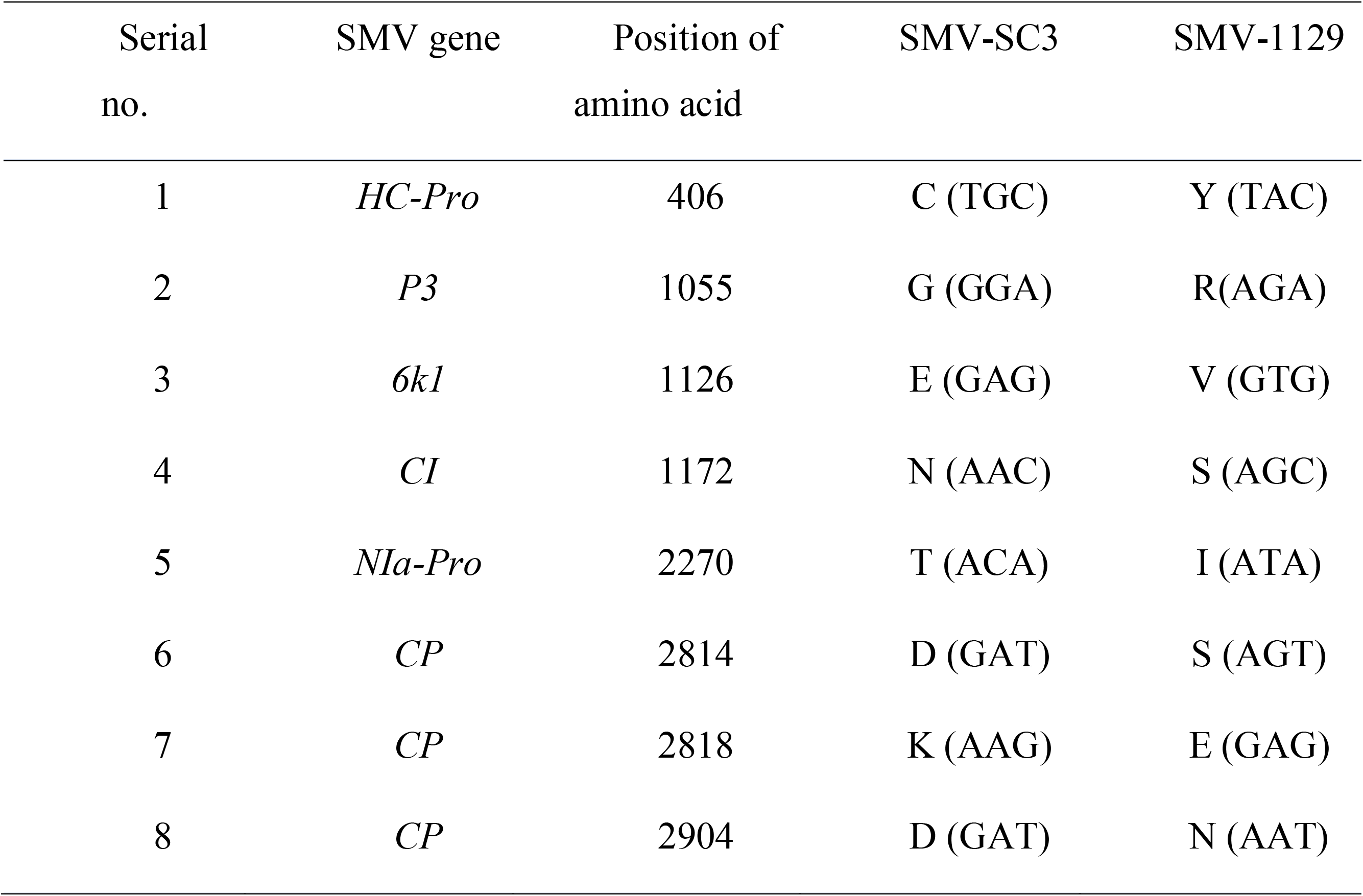
Amino acid and nucleotide differences between SMV-SC3 and SMV-1129.

The Nannong 1138-2 plants inoculated with all three recombinant clones showed systemic mosaic at 10 dpi (Figure 7B). Kefeng No.1 plants inoculated with pSMV-SC3-R3 showed mosaic symptoms at 10 dpi, while kept symptomless been inoculated with other two recombinant viral clones (Figure 7B). The ELISA assay also showed that the recombinant clone pSMV-SC3-R3 were successfully replicated in Kefeng No.1 plants, while the other two recombinant viral clones were not (Figure 7D).

**Figure 7.**
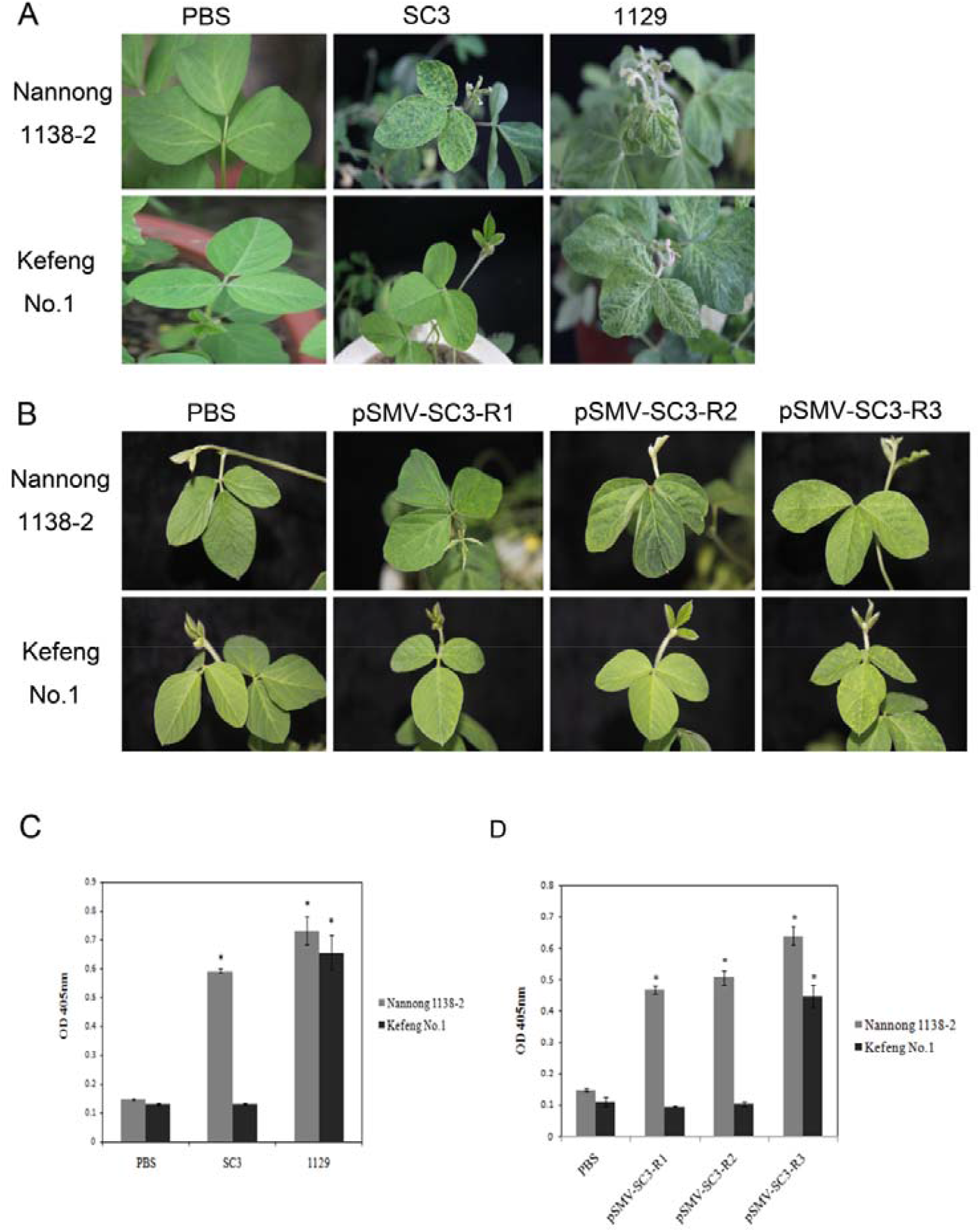
P3 or 6K1 is the virulence determinant of SMV on Kefeng No.1. Kefeng No.1 and Nannong 1138-2 plants were inoculated with PBS, SMV-SC3, SMV-1129, and recombinant virus (pSMV-SC3-R1, pSMV-SC3-R2, and pSMV-SC3-R3). Phenotypic observation and detection assay were performed at 14 dpi with SMV inoculation (A) The phenotypic responses on Kefeng No.1 and Nannong 1138-2 plants with PBS, SMV-SC3, and SMV-1129 inoculation, respectively. (B) The SMV accumulation of Kefeng No.1 and Nannong 1138-2 plants with PBS, SMV-SC3, and SMV-1129 inoculation were detected by ELISA assay. Each experiment has three replicate plants, with the standard errors indicated. Asterisks denote significant differences in the accumulation of SMV compared to that in control plants, t-test, p < 0.001. (C) The phenotypic responses on Kefeng No.1 and Nannong 1138-2 plants with PBS, pSMV-SC3-R1, pSMV-SC3-R2, and pSMV-SC3-R3 inoculation, respectively. (D) The SMV accumulation of Kefeng No.1 and Nannong 1138-2 plants with PBS, pSMV-SC3-R1, pSMV-SC3-R2, and pSMV-SC3-R3 inoculation were detected by ELISA assay. Each experiment has three replicate plants, with the standard errors indicated. Asterisks denote significant differences in the accumulation of SMV compared to that in control plants, t-test, p < 0.001.

Only two genes (P3 and 6K1) differ between the clones pSMV-SC3-R2 and pSMV-SC3-R3. The difference is located at 1055aa in P3 and 1126aa in 6K1 (Table 5, Figure 6A). Since SMV-SC3 is unable to infect Kefeng No.1 plants, we wondered whether mutations on the 1055aa or 1126aa having the corresponding amino acids from the SMV-1129 could break the resistance.

Two mutant viral clones containing either G1055R or E1126V mutations were constructed (pSMV-SC3_G1055R_ or pSMV-SC3_E1126V_) (Figure 6B) and were able to infect Nannong 1138-2 plants (Figure 8A). However, only the G1055R mutation on P3 could cause SMVSC3 to infect Kefeng No.1 plants, as shown by apparent mosaic symptoms and ELISA assay (Figure 8A and 8C).

**Figure 8.**
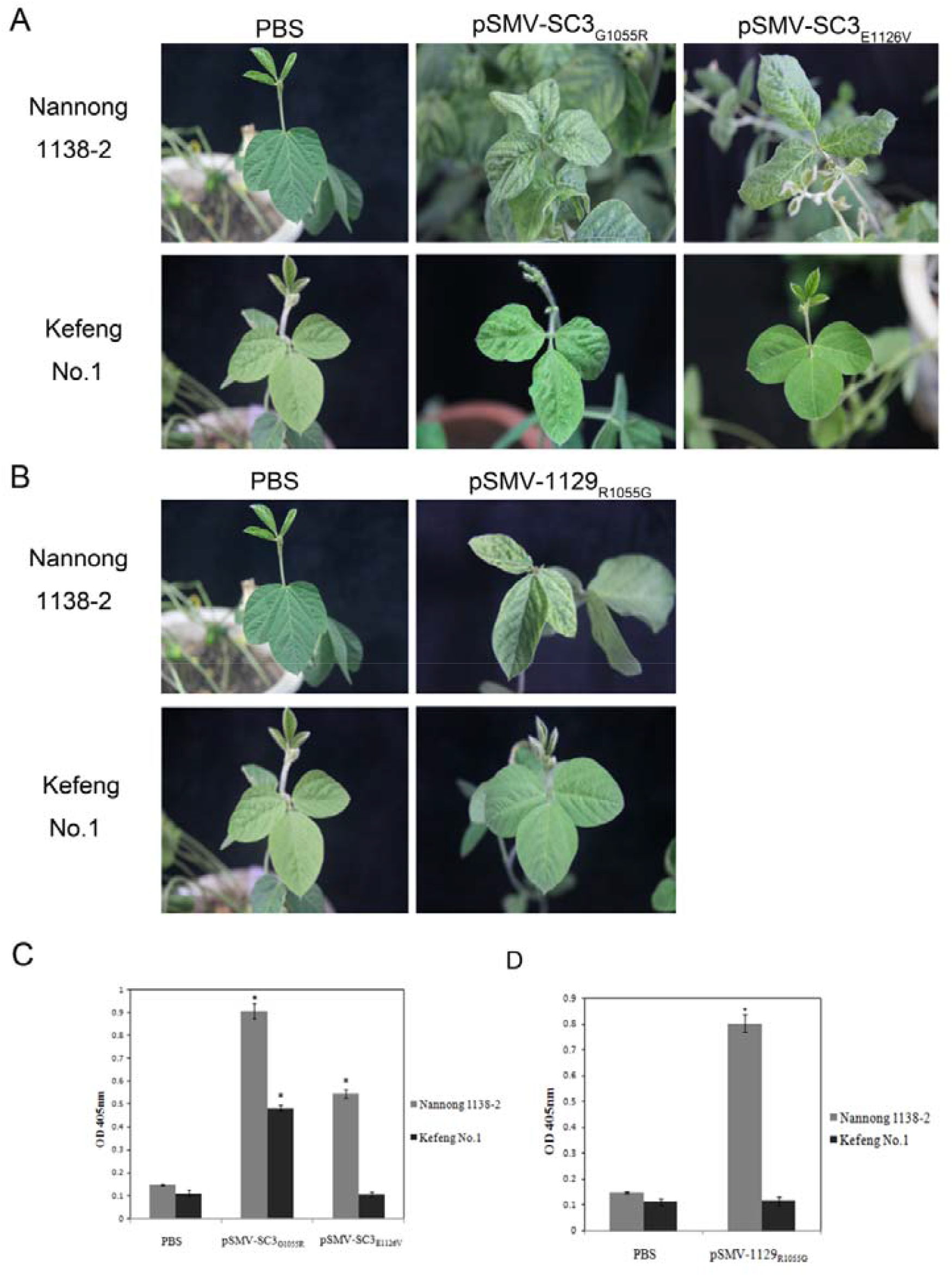
P3 determines the virulence of SMV on Kefeng No.1. Kefeng No.1 and Nannong 1138-2 plants were inoculated with PBS and point mutagenesis (pSMV-SC3_G1055R_, pSMV-SC3_E11126V_, and pSMV-1129_R1055G_). Phenotypic observation and detection assay were performed at 14 dpi with SMV inoculation. (A) The phenotypic responses on Kefeng No.1 and Nannong 1138-2 plants with PBS, pSMV-SC3_G1055R_ and pSMV-SC3_E11126V_ inoculation, respectively. (B) The SMV accumulation of Kefeng No.1 and Nannong 1138-2 plants with PBS, PBS, pSMV-SC3_G1055R_ and pSMV-SC3_E11126V_ inoculation were detected by ELISA assay. Each experiment has three replicate plants, with the standard errors indicated. Asterisks denote significant differences in the accumulation of SMV compared to that in control plants, t-test, p < 0.001. (C) The phenotypic responses on Kefeng No.1 and Nannong 1138-2 plants with PBS, and pSMV-1129_R1055G_ inoculation. (D) The SMV accumulation of Kefeng No.1 and Nannong 1138-2 plants with PBS, and pSMV-1129_R1055G_ inoculation were detected by ELISA assay. Each experiment has three replicate plants, with the standard errors indicated. Asterisks denote significant differences in the accumulation of SMV compared to that in control plants, t-test, p < 0.001.

Similarly, an R1055G mutation (pSMV-1129_R1055G_) on SMV-1129 (Figure 6B) disabled the viral ability to infect Kefeng No.1 plants, but not the Nannong 1138-2 plants (Figure 8B and 8D). These data collectively demonstrate that the P3 is the sole virulence determinant of SMV-SC3 on Kefeng No.1.

### Confirm the interaction between Rsc3K and SMV P3 via yeast two-hybrid (Y2H) assay

Since *Rsc3K* is required for the resistance of soybean cv. Kefeng No.1 to SC3, and P3 is the virulence determinant on Kefeng No.1, we decided to test the protein interaction between Rsc3K and SMV P3 by using Y2H assay. Yeast cells were transformed with plasmid to express Rsc3K, together with another plasmid to express SMV-SC3 P3, or SMV-1129 P3, or an empty plasmid as a negative control. Yeast cells transformed with plasmids for the expression of SMV-SC4 CP and GmCPIP were used as a positive control (Zong *et al*., 2020). Sequentially diluted yeast cells were spotted onto double dropout and quadruple dropout media (Figure 9A and 9B). All cells were grown on the double dropout medium, while the positive control or cells expressing Rsc3K and P3 from both SMV strains can grow on the quadruple dropoutmedium (Figure 9B). Interestingly, we found that cells expressing Rsc3K and SMV-SC3 P3 had grown more colonies than that of the cells expressing Rsc3K and SMV-1129 P3 at the same initial concentration. Anyway, the Y2H experiments indicated protein interaction between Rsc3K and SMV P3.

**Figure 9.**
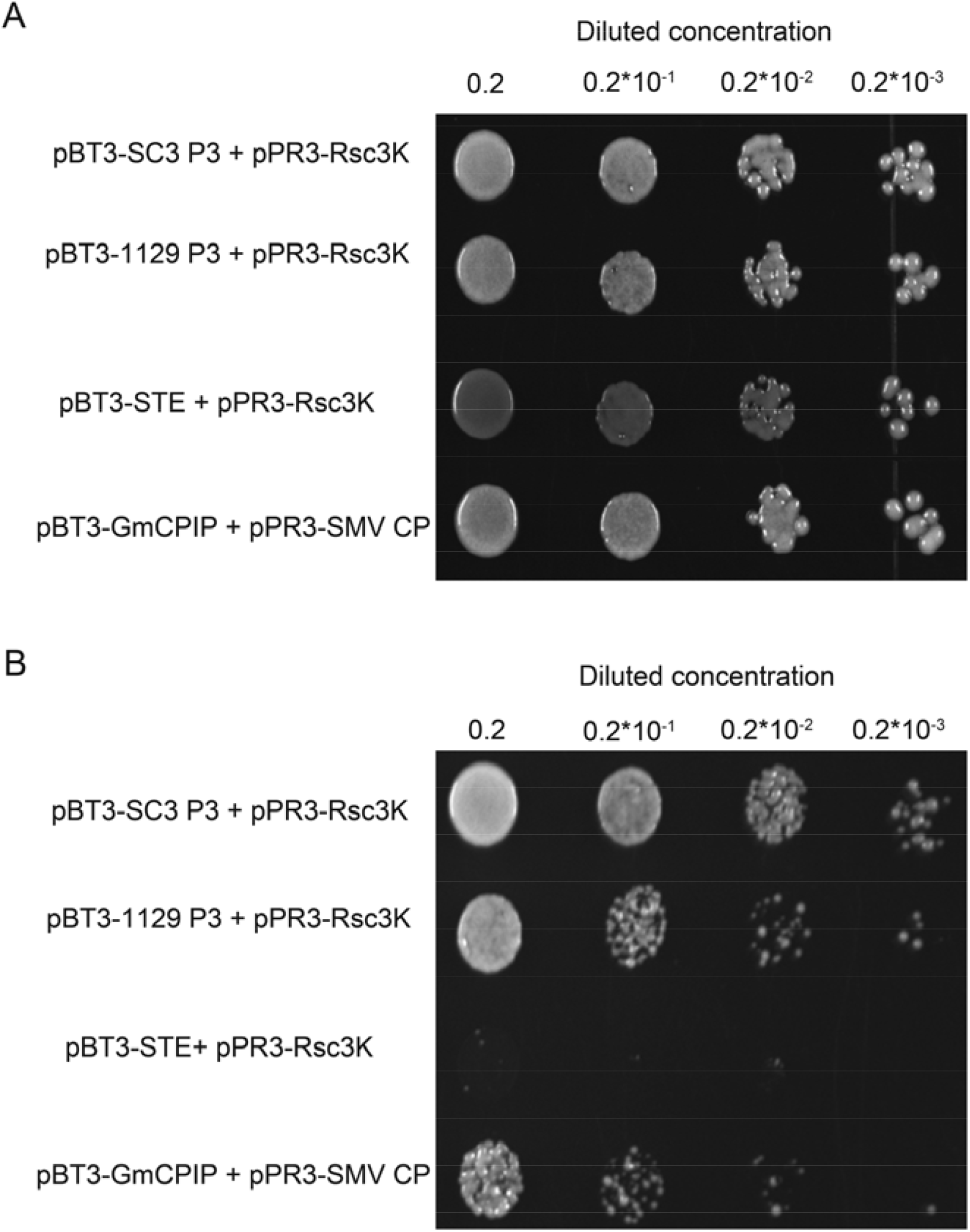
Interaction between Rsc3K and SMV P3 (SC3 and 1129) were determined in the yeast two-hybrid system. Yeast cells transformed with plasmids for the expression of SMV-SC4 CP and GmCPIP were used as the positive control. Coexpression of pBT3-STE and pPR3-N was the negative control. Sequentially diluted yeast cells were spotted onto double dropout and quadruple dropout media. The initial concentration of yeast in the medium is 0.2 (OD_600_=0.2). (A) Yeast cells transformed with bait (pBT3-SC3 P3 and pBT3-1129 P3) and prey (pPR3-Rsc3K) vectors were grown on double dropout medium to assess their double transformation. Images were taken at 2 days post transformed. (B) Yeast cells transformed with bait (pBT3-SC3 P3 and pBT3-1129 P3) and prey (pPR3-Rsc3K) vectors were grown on double dropout medium to assess their double transformation. Images were taken at 4 days post transformed.

### Interaction verification experiment between Rsc3K and SMV P3 by bimolecular fluorescent complementation (BiFC)

The BiFC assay was used to further confirm the interaction between Rsc3K and SMV P3 in agrobacterium-infiltrated *N. benthamiana* plants. Rsc3K was fused with the N-terminal YFP fragment (Rsc3K -NYFP), and SMV P3 was fused with the C-terminal YFP fragment (SMV P3-CYFP). Co-expression of Rsc3K-NYFP and SMV SC3 P3-CYFP induced strong YFP fluorescence signals in agroinfiltrated *N.benthamiana* leaf cells under a confocal microscope (Figure 10). As controls, YFP fluorescence was not observed in *N. benthamiana* leaf cells expressing Rsc3K-NYFP + CYFP or NYFP + SMV P3-CYFP (Figure 10). These results confirmed that Rsc3K interacted with SMV P3. In addition, we found the fluorescence intensity of Rsc3K-NYFP and SMV 1129 P3-CYFP did not increase significantly even when the *Agrobacterium* concentration was increased. These results suggested that Rsc3K and SMV-SC3 P3 may have stronger binding affinity than that of Rsc3K and SMV-1129.

**Figure 10.**
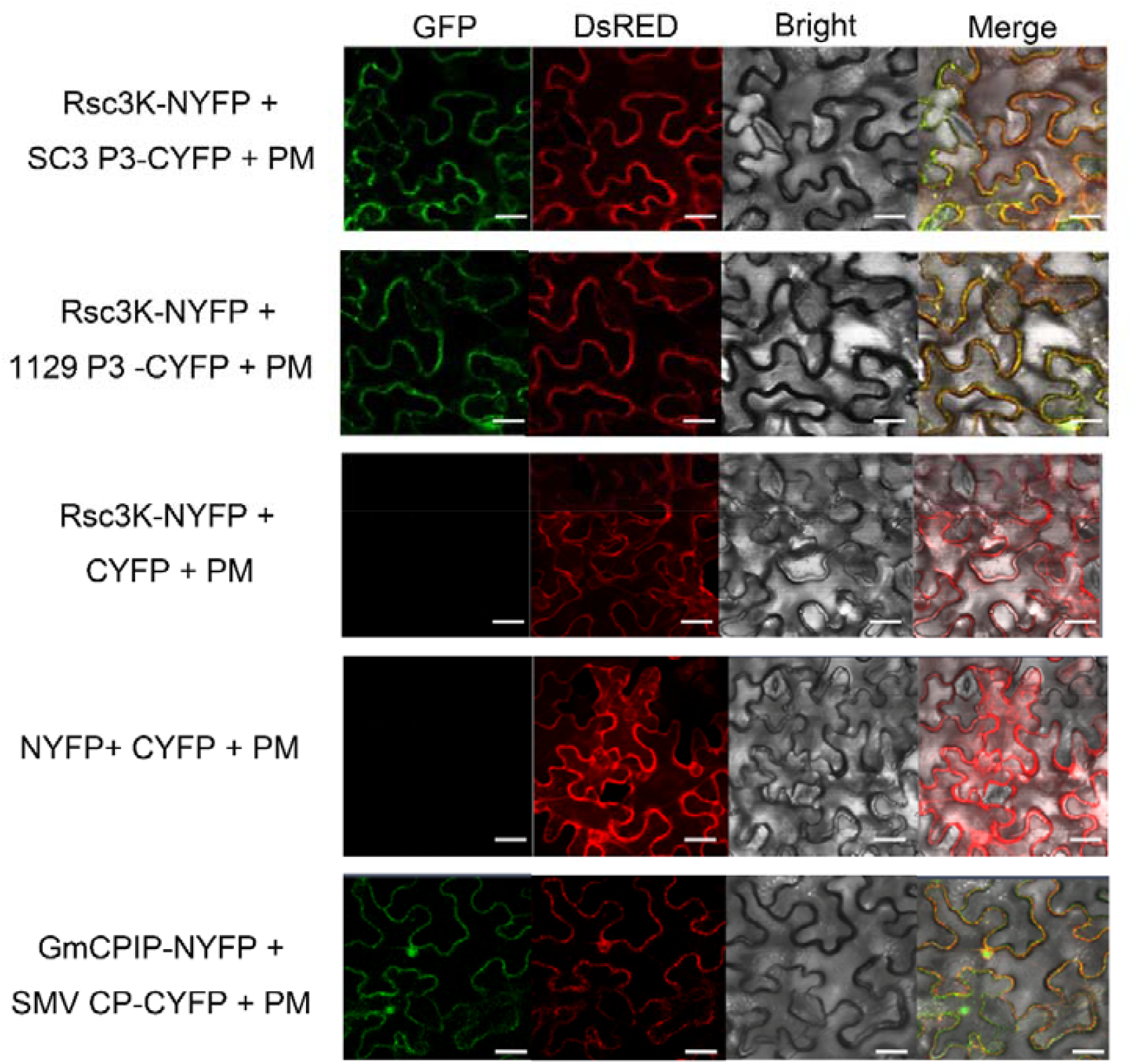
Interaction between Rsc3K and SMV P3 (SC3 and 1129) were confirmed by bimolecular fluorescent complementation assay. The expression vectors Rsc3K-NYFP and SMV P3-CYFP (SC3 P3-CYFP and 1129 P3-CYFP) with PM marker were transmitted into *N. benthamiana* epidermal leaves. Coexpression of Rsc3K-NYFP and CYFP, or NYFP and CYFP, was the negative control, GmCPIP-NYFP and SMV CP-CYFP, was the positive control. The fluorescence signal was observed at 4 days postinoculation. GFP, RFP, bright, and merged fluorescence images are shown. Scale bars represent 10 μm.

### Two RNase H family proteins of susceptible cultivars also interact with SMV P3

Previous experimental results showed that *Rsc3K* in Kefeng No.1 (resistant) genome was produced by deletion and recombination of two genes (*LOC100526921* and *LOC100812666*) in Williams 82 (susceptible) genome. Through gene model analysis, we found that these three genes (*Rsc3K* , *LOC100526921*, and *LOC100812666*) encode RNase proteins. Since the domains of the three genes are so similar, why could the recombinant *Rsc3K* mediate the resistance of Kefeng No.1 to SMV-SC3, while Williams 82 carrying *LOC100526921* and *LOC100812666* shows susceptibility to SMV-SC3? Since Rsc3K interact with SMV virulence determinant P3, we speculate that this interaction may be related to the resistance mechanism. So are these two proteins (LOC100526921 and LOC100812666) unable to interact with P3, resulting in Williams’ sensitive symptoms? In order to verify this hypothesis, we used Y2H and BiFC experiments to verify the interaction between the proteins LOC100526921 and LOC100812666 and SMV P3. The experimental results show that the proteins encoded by *LOC100526921* and *LOC100812666* also interact with SMV P3 (Figure 11 and 12). Therefore, the susceptibility of Williams 82 to SMV may not be due to the loss of the ability to interact with the virulence determinant P3.

**Figure 11.**
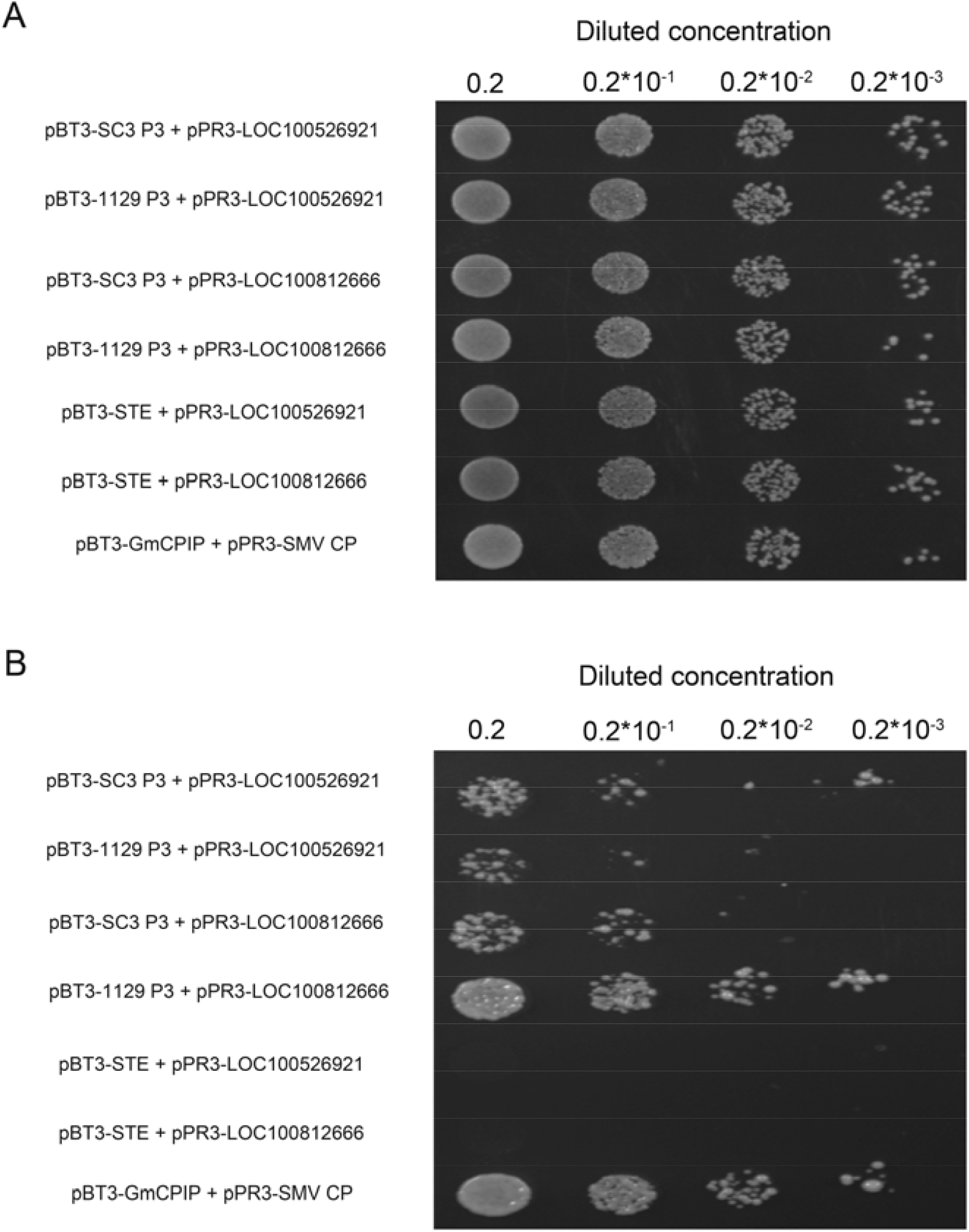
Interaction between LOC100516921 or LOC100812666 and SMV P3 (SC3 and 1129) were determined in the yeast two-hybrid system. Yeast cells transformed with plasmids for the expression of SMV-SC4 CP and GmCPIP were used as the positive control. Coexpression of pBT3-STE and pPR3-LOC100516921 or pPR3-LOC100812666, were negative controls. Sequentially diluted yeast cells were spotted onto double dropout and quadruple dropout media. The initial concentration of yeast in the medium is 0.2 (OD_600_=0.2). (A) Yeast cells transformed with bait (pBT3-SC3 P3 and pBT3-1129 P3) and prey (pPR3-LOC100516921 and pPR3-LOC100812666) vectors were grown on double dropout medium to assess their double transformation. Images were taken at 2 days post transformed. (B) Yeast cells transformed with bait (pBT3-SC3 P3 and pBT3-1129 P3) and prey (pPR3-LOC100516921 and pPR3-LOC100812666) vectors were grown on double dropout medium to assess their double transformation. Images were taken at 4 days post transformed.

**Figure 12.**
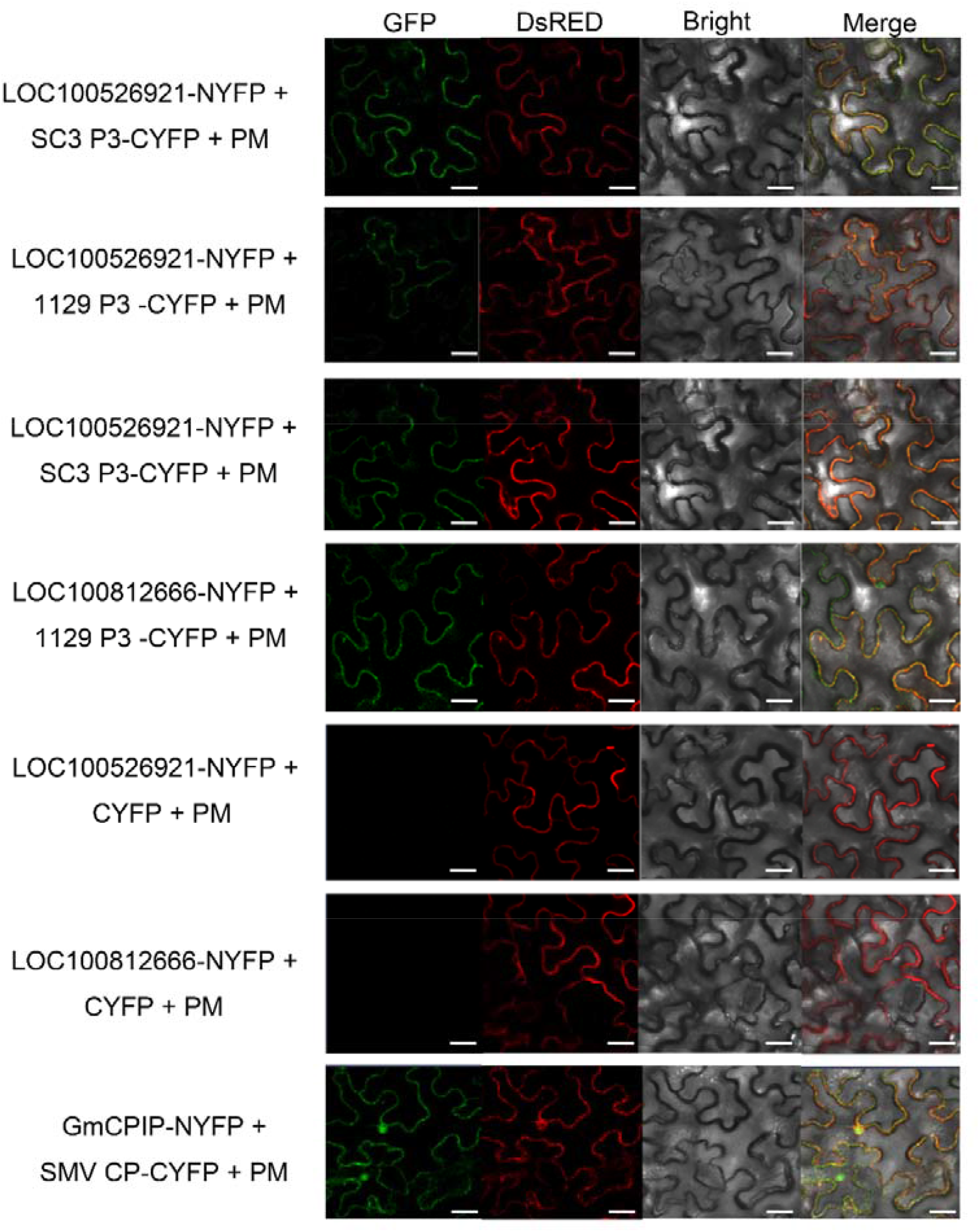
Interaction between LOC100516921 or LOC100812666 and SMV P3 (SC3 and 1129) were confirmed by bimolecular fluorescent complementation assay. The expression vectors LOC100516921-NYFP and SMV P3-CYFP (SC3 P3-CYFP and 1129 P3-CYFP), or LOC100812666-NYFP and SMV P3-CYFP with PM marker were transmitted into *N. benthamiana* epidermal leaves. Coexpression of LOC100516921-NYFP and CYFP, or LOC100812666-NYFP and CYFP, was the negative control, GmCPIP-NYFP and SMV CP-CYFP, was the positive control. The fluorescence signal was observed at 4 days postinoculation. GFP, RFP, bright, and merged fluorescence images are shown. Scale bars represent 10 μm.

### Bioinformatic analysis of LOC100526921, LOC100812666, and Rsc3K

The results of Y2H and BiFC experiments showed that the two proteins (LOC100526921 and LOC100812666) also interact with P3, indicating that there are other factors that cause Williams 82 to show susceptible symptoms to SMV. Through the transmembrane structure prediction of Rsc3K, LOC100526921, and LOC100812666, we found that the three proteins all have a transmembrane structure, but the N-terminus of Rsc3K and LOC100526921 is inside the transmembrane region, and the C-terminus is outside the transmembrane region, while LOC100812666 is just the opposite (Supplementary Figure 1). By comparing the amino acid sequences of Rsc3K, LOC100526921 and LOC100812666, we found that the 208 position of LOC100526921 is aspartic (208D), which corresponds to the asparagine (216N) at 216 position of LOC100812666 and the asparagine (221N) at 221 position of Rsc3K (Supplementary Figure 2). In short, Rsc3K and LOC100812666 belong to DEDN-model dsRNAse, while LOC100526921 belongs to DEDD-model dsRNAse.

### P3 Polymorphism determines virulence

According to previous studies, the virulent determinance of SMV on *Rsv4*-genotype soybean is P3 protein, and the virulence determination sites of different SMV isolates in P3 are different. After the single nucleotide substitution (Q1033K or G1054R) in P3 of SMV-N (SMV-G2 strain), the mutant has the ability to infect V94-5152 (*Rsv4*-genotype soybean) (Chowda *et al*., 2011b; Khatabi *et al*., 2012; Wang *et al*., 2015; Ishibashi *et al*. 2019). In this study, the G1055R mutation changed the virulence of SMV-SC3 on Kefeng No.1. This mutation site is inconsistent with previous reports, but through sequence alignment, we found the polymorphic site 1055 of SMV-SC3 corresponds to the 1054 site of SMV-N, and the polymorphic site 1034 of SMV-SC3 corresponds to the 1033 site of SMV-N (Supplementary Figure 3). This indicates that the substitution of Q1033K/Q1034K or G1054R/G1055R is very important for SMV to abtain the ability to infect *Rsv4*-genotype soybean. By predicting the structure of the SMV P3 protein, we found that P3 has two transmembrane helices, the 1034 site is located in the second transmembrane helix region, and the 1055 site is located outside the transmembrane helix (Supplementary Figure 4). However, the amino acid mutation at these two sites did not lead to the change of the original structure. The structural regions of the two virulence determination sites are different, but after the corresponding amino acid substitutions, the mutants can overcome the resistance of *Rsv4*-genotype soybean, indicating that the virulence determining sites may have nothing to do with the transmembrane structure. We speculate that the amino acid replacement at the corresponding sites may affect other functions of the P3 protein (such as the ligand binding, etc.), causing the mutant to break the original disease resistance mechanism of the soybean.

Khatabi *et al*. (2012) identified the infectivity of multiple SMV isolates in North America on *Rsv4*-genotype soybeans and found that isolates with Q1033K or G1054R variants have the virulence on V94-5152 (*Rsv4*-genotype soybean). We analyzed the phylogenetic tree of P3 in Chinese SMV isolates and found that there was no Q1033K or G1054R variation in other isolates except SMV-1129 (with G1054R variation). This indicates that the G1055R mutation of SMV may have occurred independently in China (Figure 13). This means that the disease resistance mechanism of *Rsv4*-genotype soybeans is beginning to face challenges, prompting us to accelerate the discovery of other SMV resistance genes and pay more attention to polymerization breeding.

**Figure 13.**
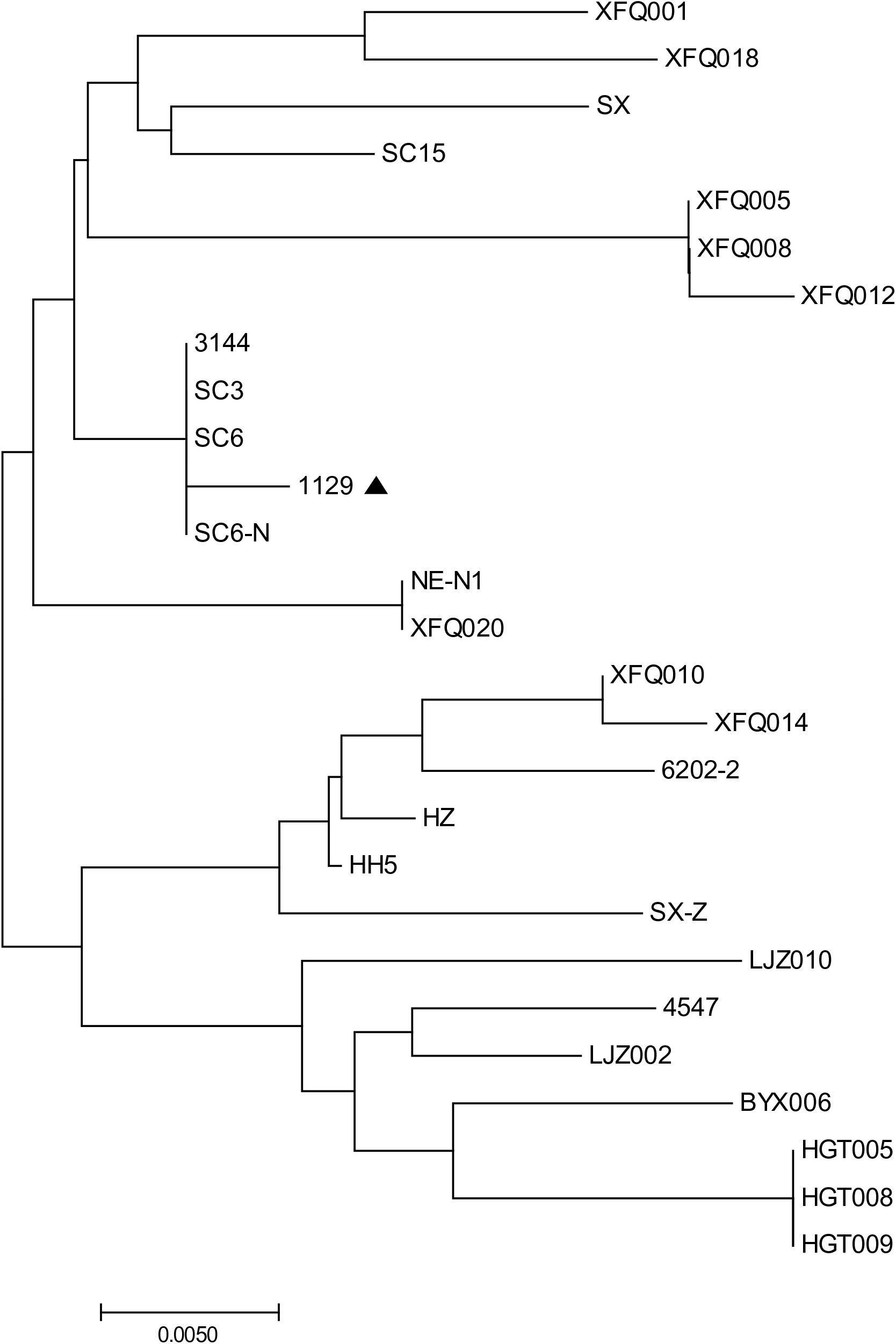
Phylogenetic tree analysis of P3 proteins of Chinese SMV isolates. Phylogenetic tree of SMV P3 proteins is constructed by adopting the Neighbor-Joining method. The optimal tree with the sum of branch length = 0.17203854 is shown. The triangle represents the virulence determining site of SMV P3 protein is G1054R/G1055R. The Genbank accession number of SMV isolates used for evolutionary analyses are listed in Supplemental Table 2.

## Discussion

SMV-SC3 is one of the prevalent strains in China’s Huang-Huai and Changjiang Valleys regions, but the resistance gene of soybean to SMV-SC3 has not been cloned so far (Guo *et al*., 2005; Li *et al*., 2010; Zheng *et al*., 2014). In this study, the F_2:3_ and RIL populations crossed by Kefeng No.1 and Nannong 1138-2 were used to locate the resistance gene of SMV-SC3. It was found that the resistance locus was located on chromosome 2 and named *Rsc3k*. Interestingly, there were many resistance loci of Kefeng No.1 to other SMV strains near the resistance locus of *Rsc3K*. The localization intervals of these resistance loci overlap with each other (Supplementary Figure 5). However, no resistance gene has been identified. Through VIGS experiment, we found that the *Rsc3K*-silenced plants all lost resistance to SMV strains (SMV-SC3, SMV-SC5, SMV-SC7, SMV-SC8, SMV-SC10, and SMV-SC18), which implies that the resistance of Kefeng No.1 to SMV multiple strains may be mediated by *Rsc3K*. Recently, Ishibashi *et al*. (2019) reported that *Rsv4*, a resistance gene from soybean cv. Peking, encodes an RNase H family protein, which degrades the dsRNA of the virus by interacting with the P3 of SMV to enter the virus replication compartment, and then mediates Peking’s resistance to SMV. Interestingly, *Rsv4* is a recombinant gene resulting from a 3.6 kb deletion in Peking’s genome. Through sequence alignment, we found that Rsv4 is 100% identical in amino acids level to Rsc3k, suggesting that the two genes have the same function.

The results of protein secondary structure prediction showed that the transmembrane structure of LOC100812666 of susceptible cultivar Williams 82 was different from that of resistant cultivar Kefeng No.1 (Supplementary Figure 1). We speculate that different transmembrane structures may cause differences in the perception of pathogens by Rsc3K/LOC100812666 or the interaction between Rsc3K/LOC100812666 and other genes in the resistance pathway. Therefore, *LOC100812666* may not mediate the resistance of Williams 82 to SMV strains. According to previous studies, the DEDD model of RNase H protein can degrade the RNA bands of DNA-RNA hybrids, while the DEDN model restores the activity of degrading the RNA bands of DNA-RNA hybrids under the catalysis of manganese ions (Nowotny *et al*., 2005). And the Rsv4 with DEDN model has the activity of degrading dsRNA under the catalysis of manganese ion (Ishibashi *et al*., 2019). Therefore, we speculate that LOC100526921 with DEDD model (Supplementary Figure 2) may not have strong dsRNA degradation activity, or Williams 82 plants may not have sufficient conditions (such as manganese ion deployment) to make LOC100526921 completely degrade the dsRNA of the virus, resulting in Williams 82 becoming susceptible to SMV strains. These conjectures need more experiments to verify.

In conclusion, we have finely mapped and verified that *Rsc3K* is the gene that mediates the resistance of Kefeng No.1 to SMV. Furthermore, *Rsc3K* is a recombinant gene formed due to genome deletion, which may mediates resistance to SMV by degrading viral dsRNA. The results can be used for molecular resistance breeding and tracking the evolutionary history of soybeans.

## Author Contributions

Conceptualization, Z.H. and X.K.; Methodology, Z.H. and W.L.; Investigation, J.T., Y.J., X.S., Y.Y., L. H., and L. M.; Resources, Z.H., W.L., L.B., Z.T., and L.K.; Writing, Z.H., and J.T.; Supervision, Z.H., and W.L.; Funding Acquisition, Z.H. and W.L.

## Acknowledgments

We thank John H. Hill (Iowa State University), and Zhenghe Li (Zhejiang University) for offering pBPMV, pCB301-CEN, and yeast strain W303-1B as gifts. This work was supported by the Fundamental Research Funds for the Central Universities (KYT201801), Program for Changjiang Scholars and Innovative Research Team in University (PCSIRT_17R55), the National Soybean Industrial Technology System of China (CARS-004), and Jiangsu Collaborative Innovation Center for Modern Crop Production (JCIC-MCP).

## Data availability

The data supporting the findings of this study are included within the article and its supplementary data published online.

## Supplemental data

**Supplemental Table 1.**
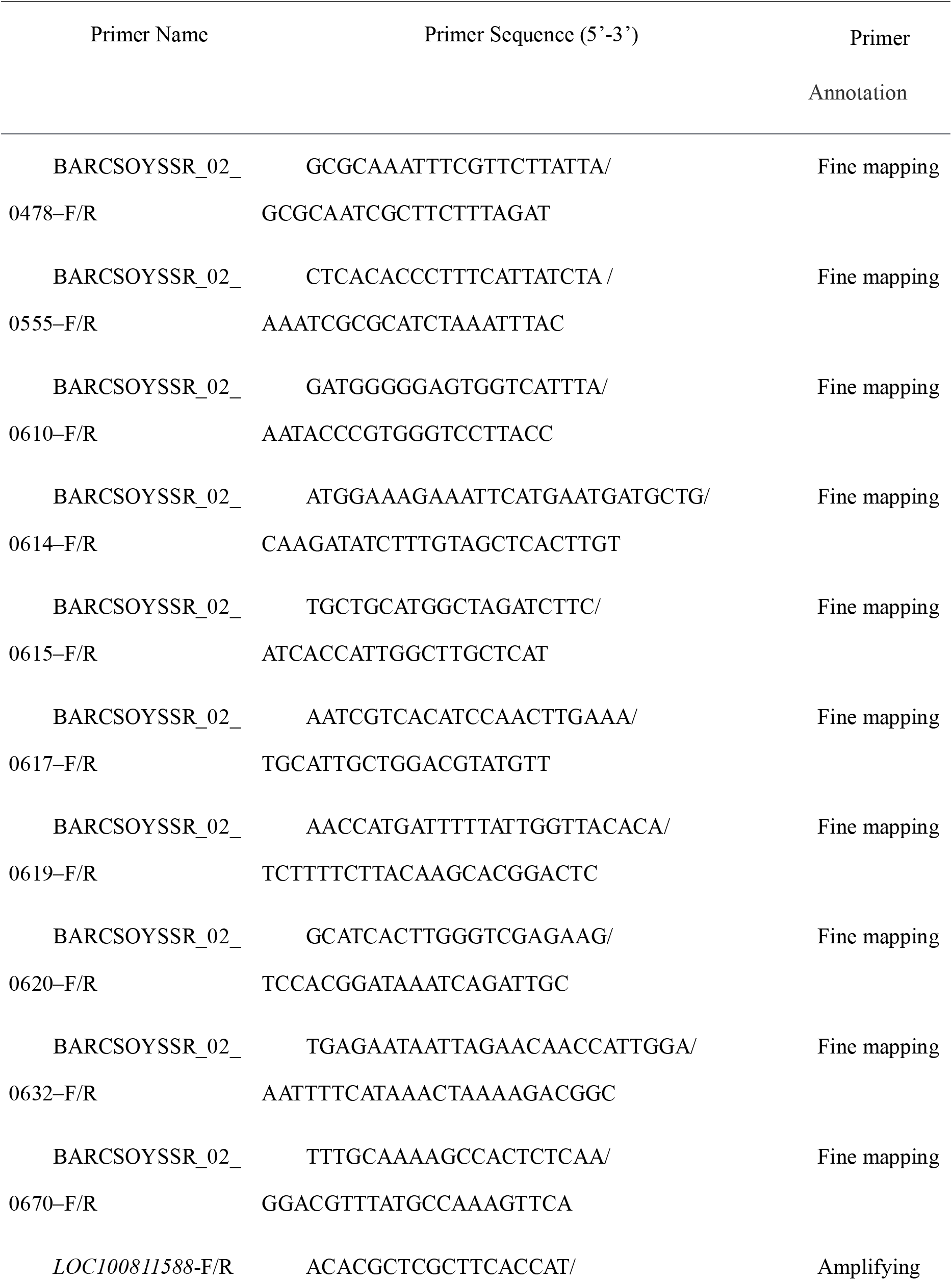
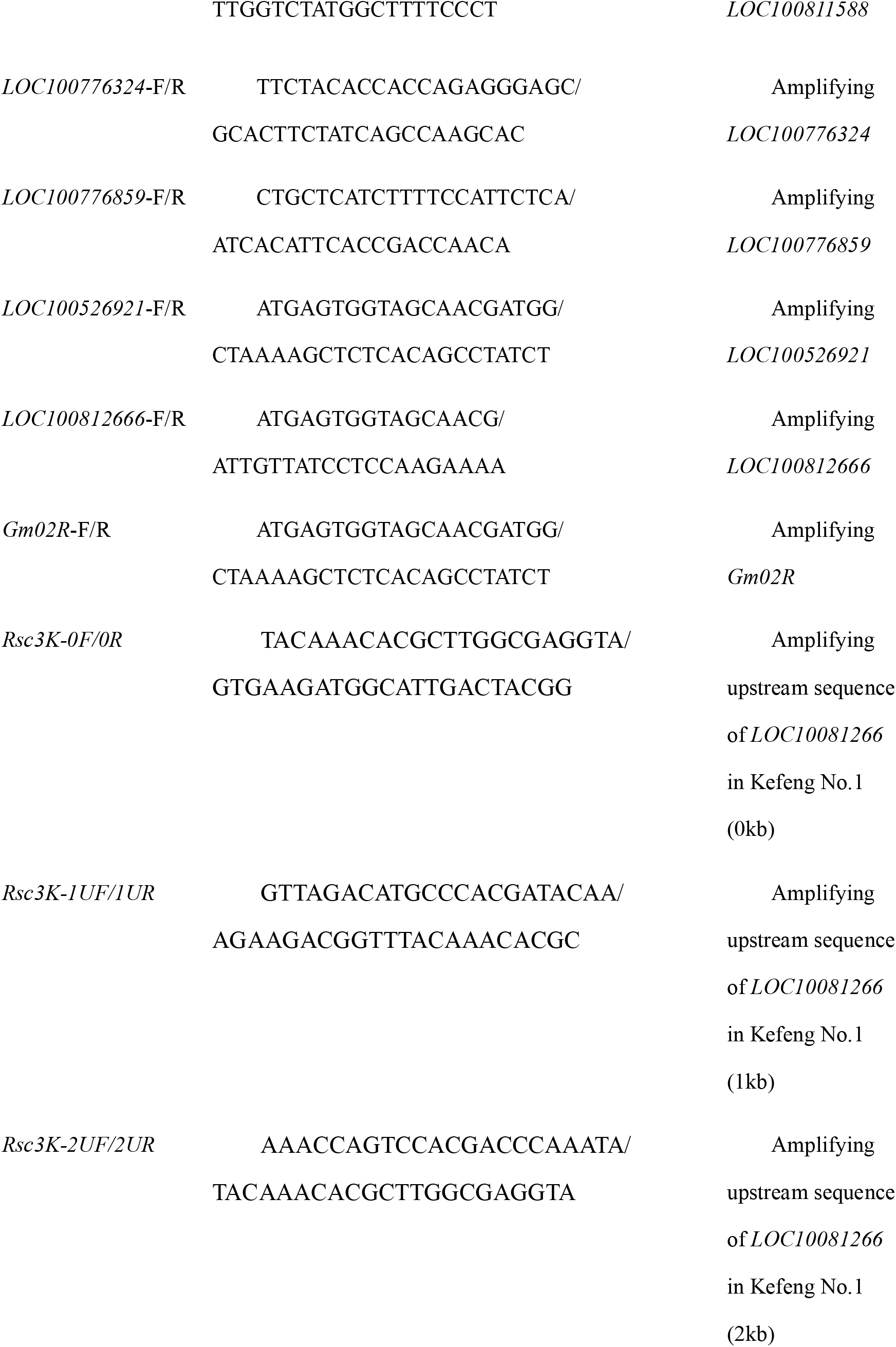
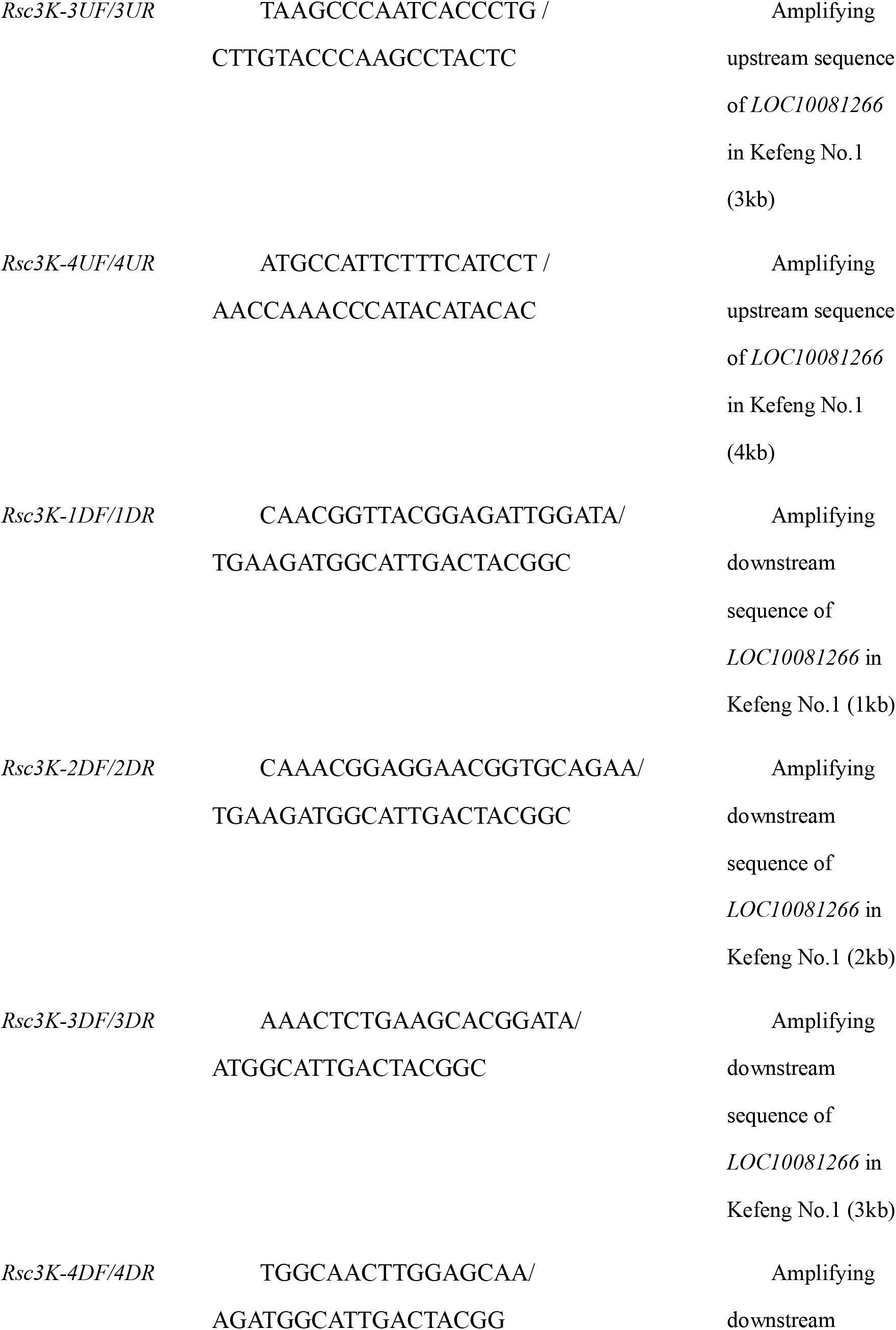
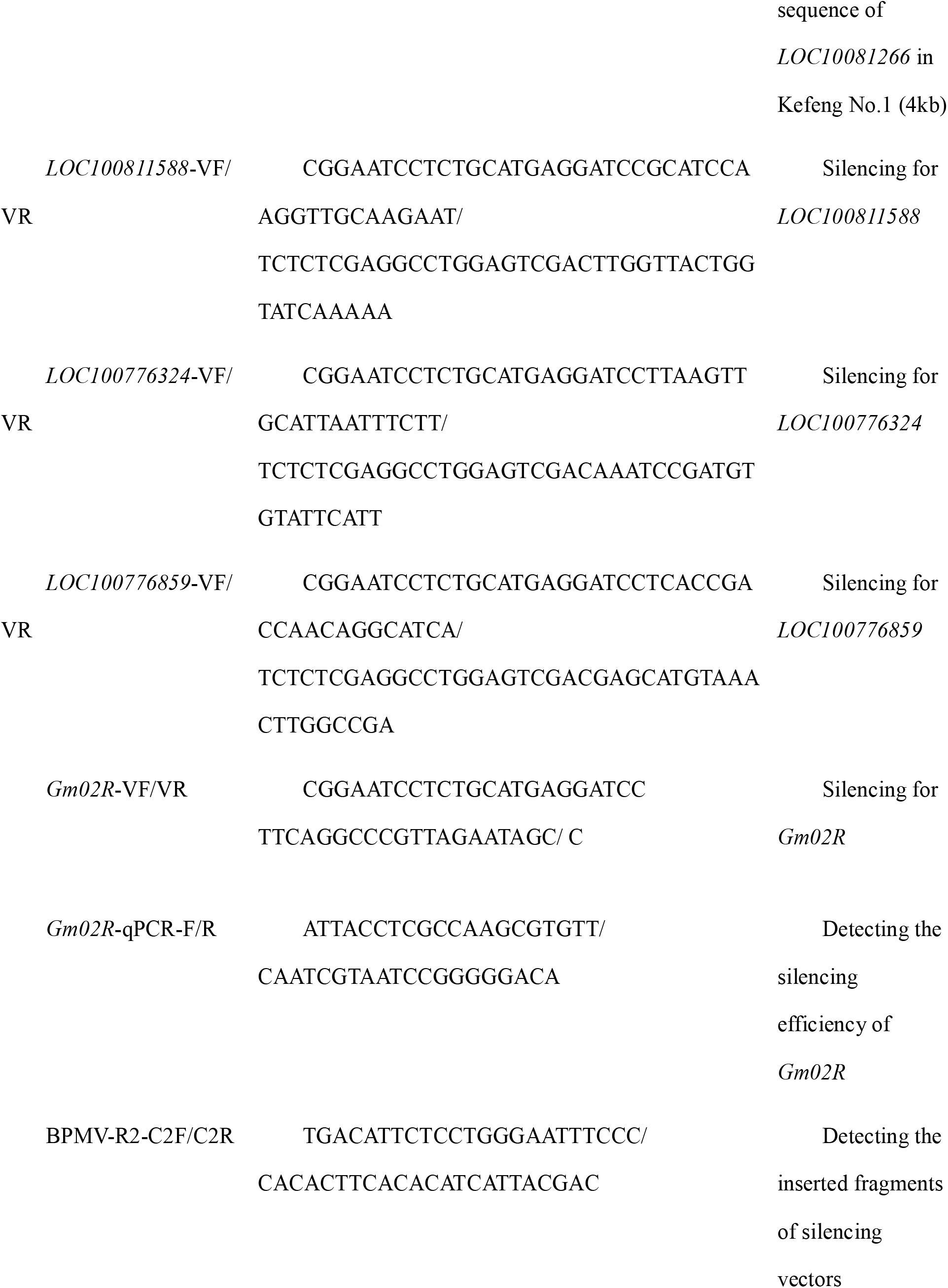
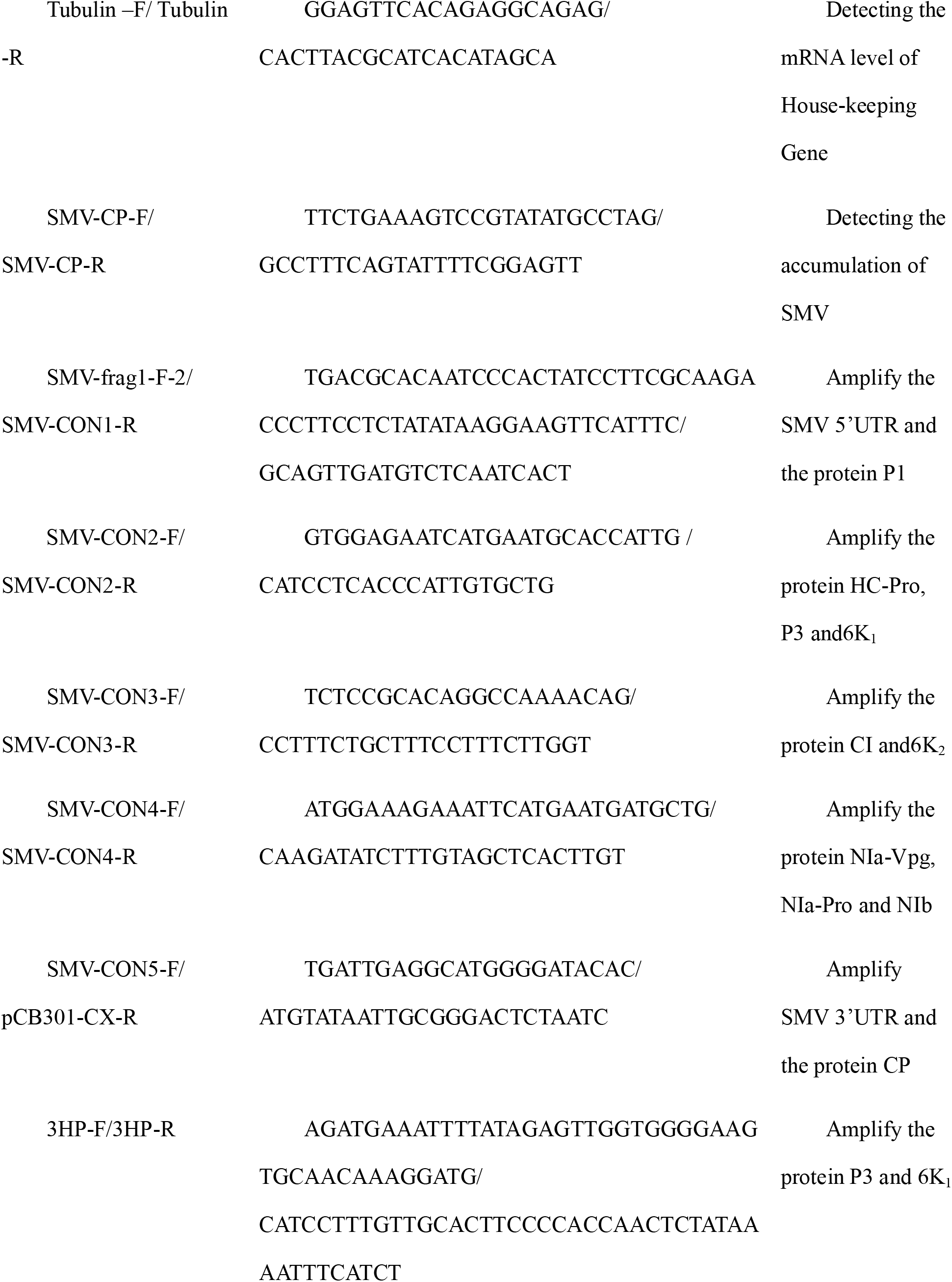
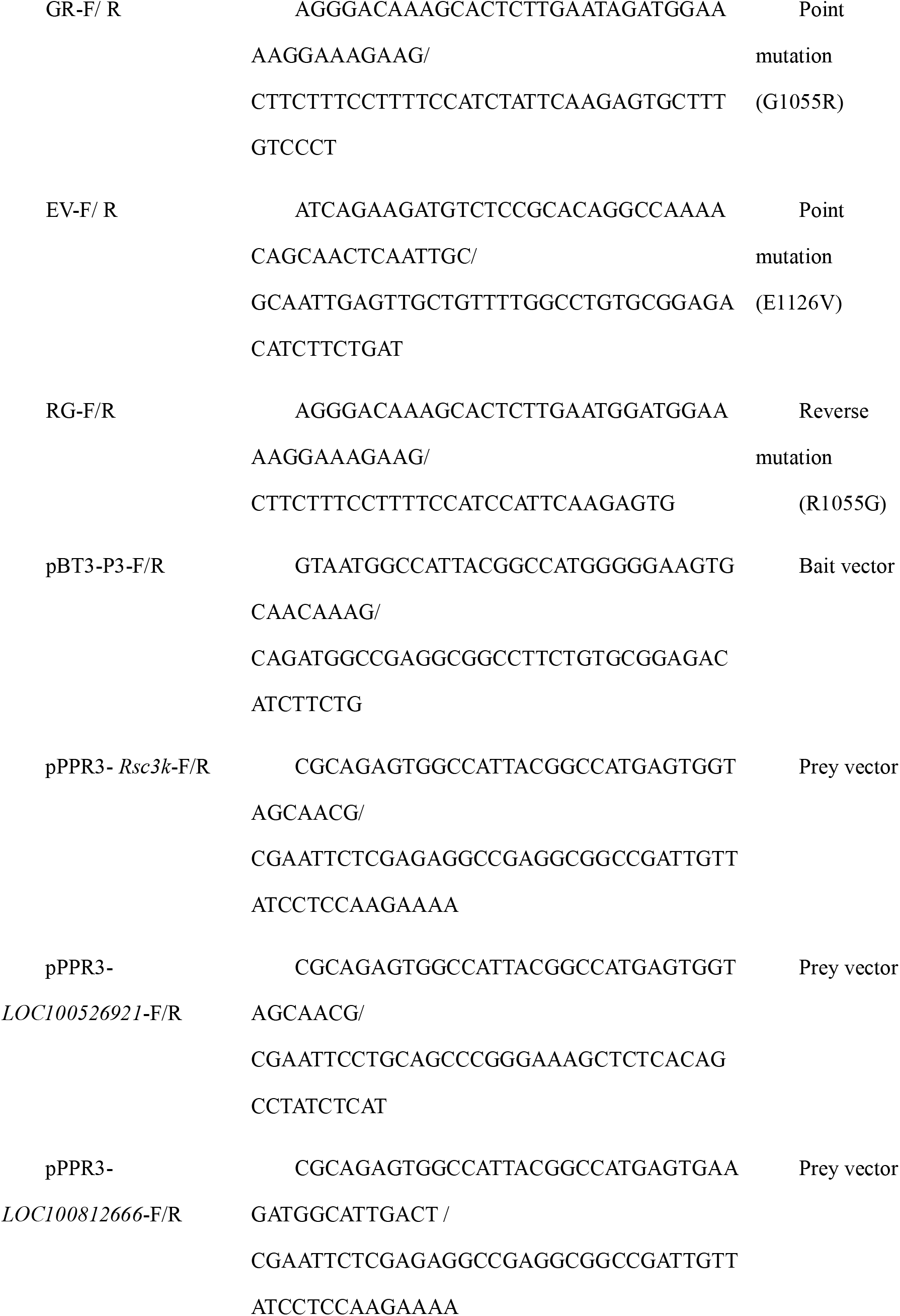
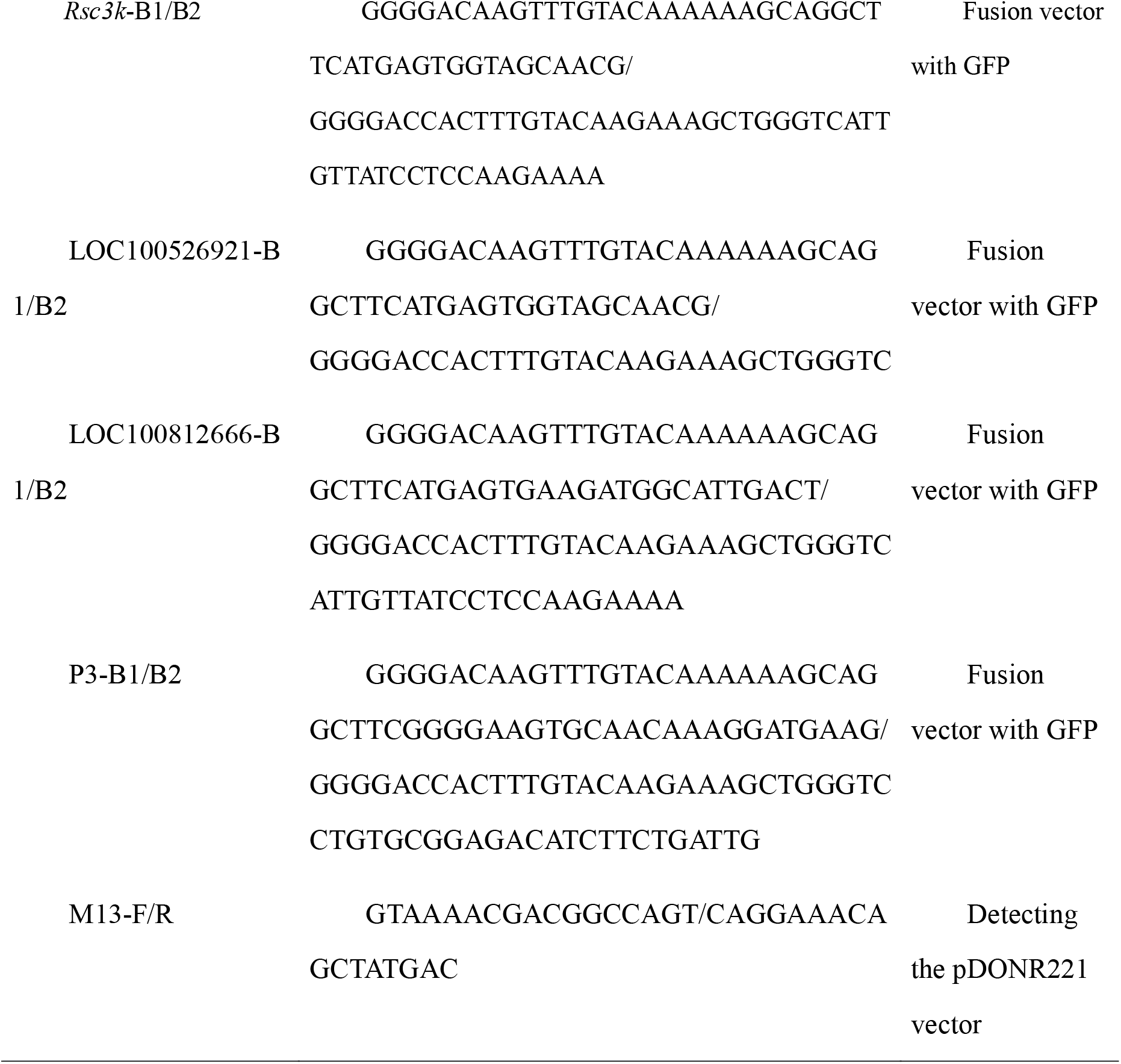
Primers used in this study.

**Supplemental Table 2.**
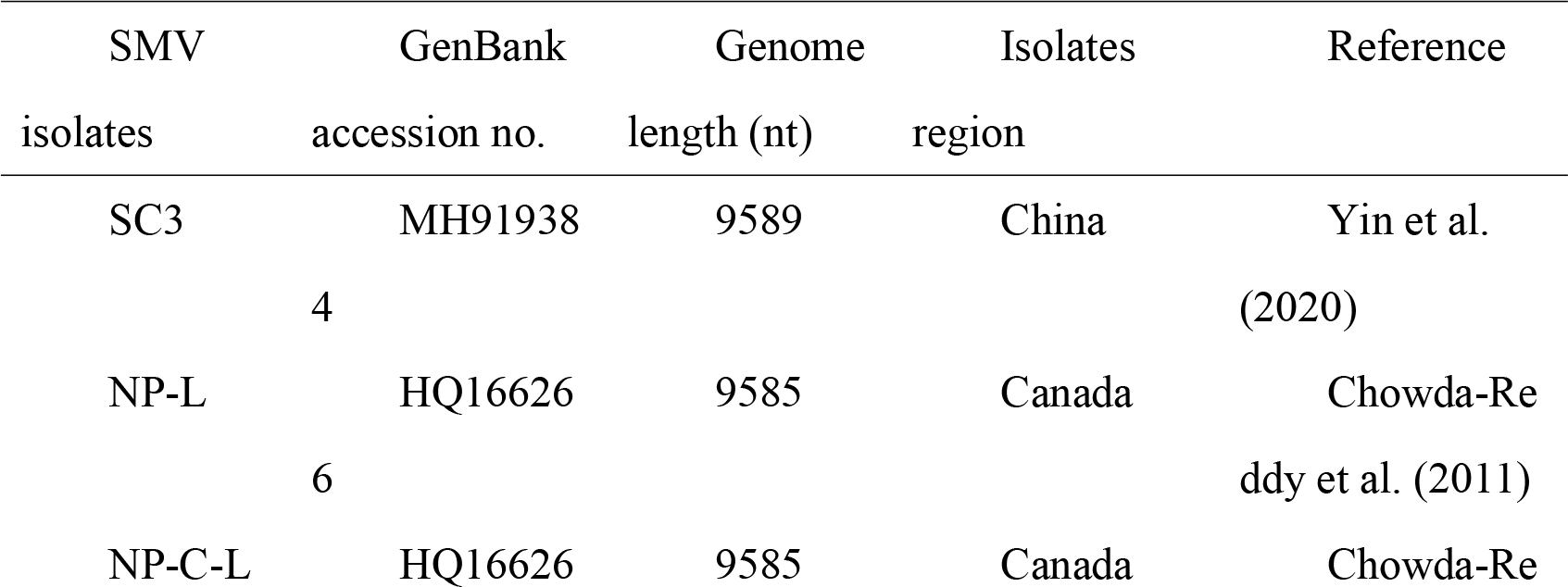
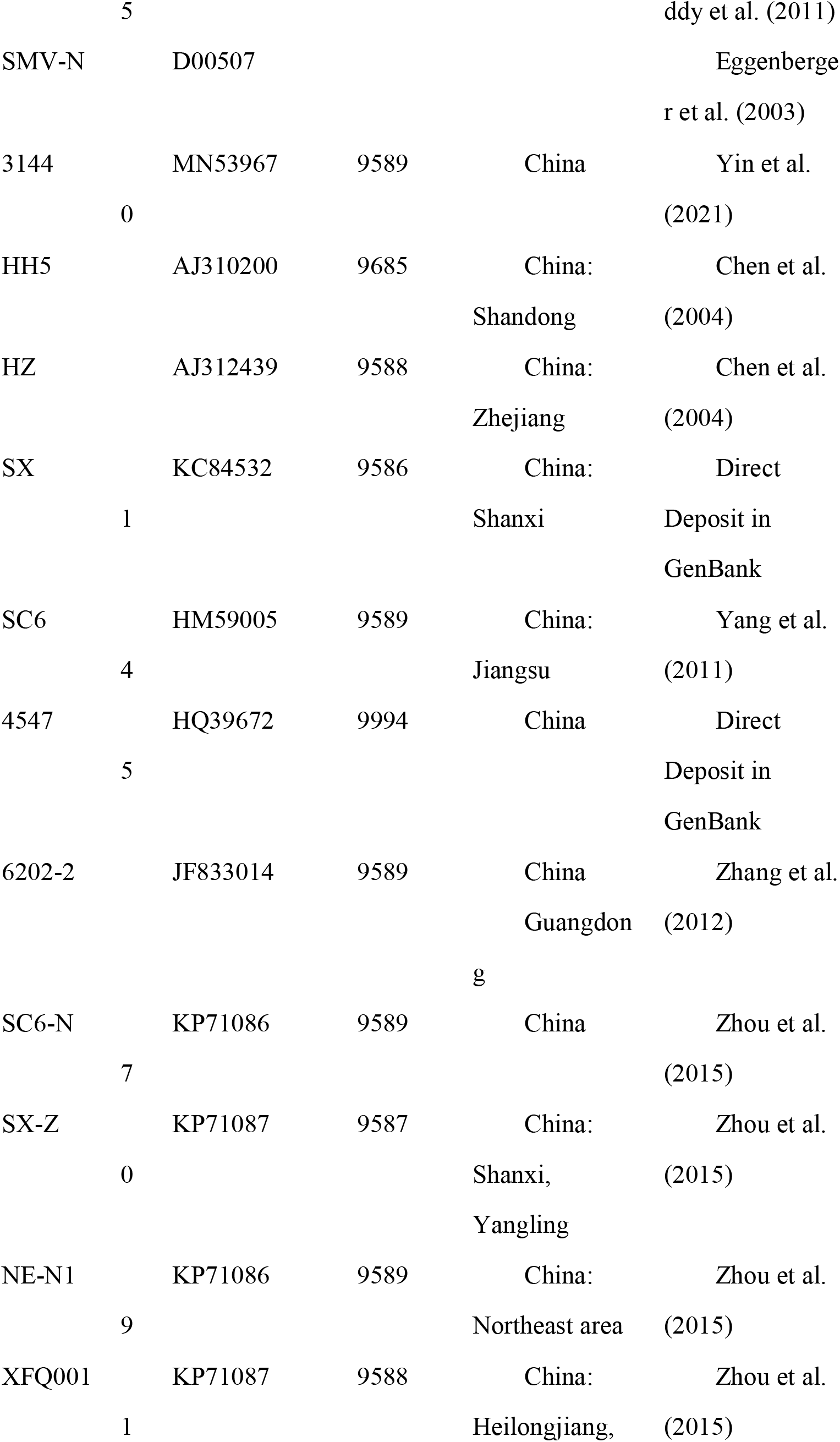
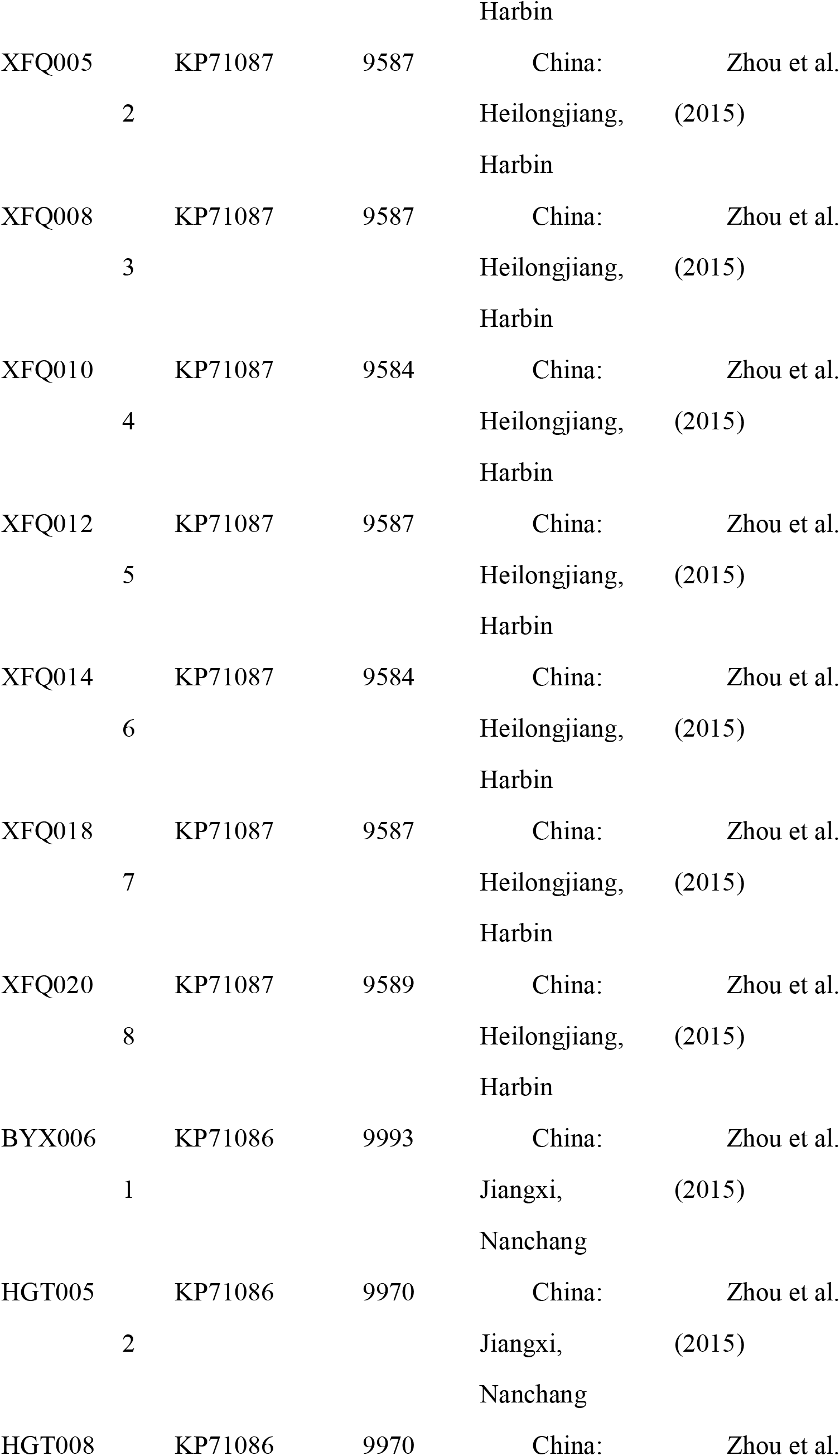
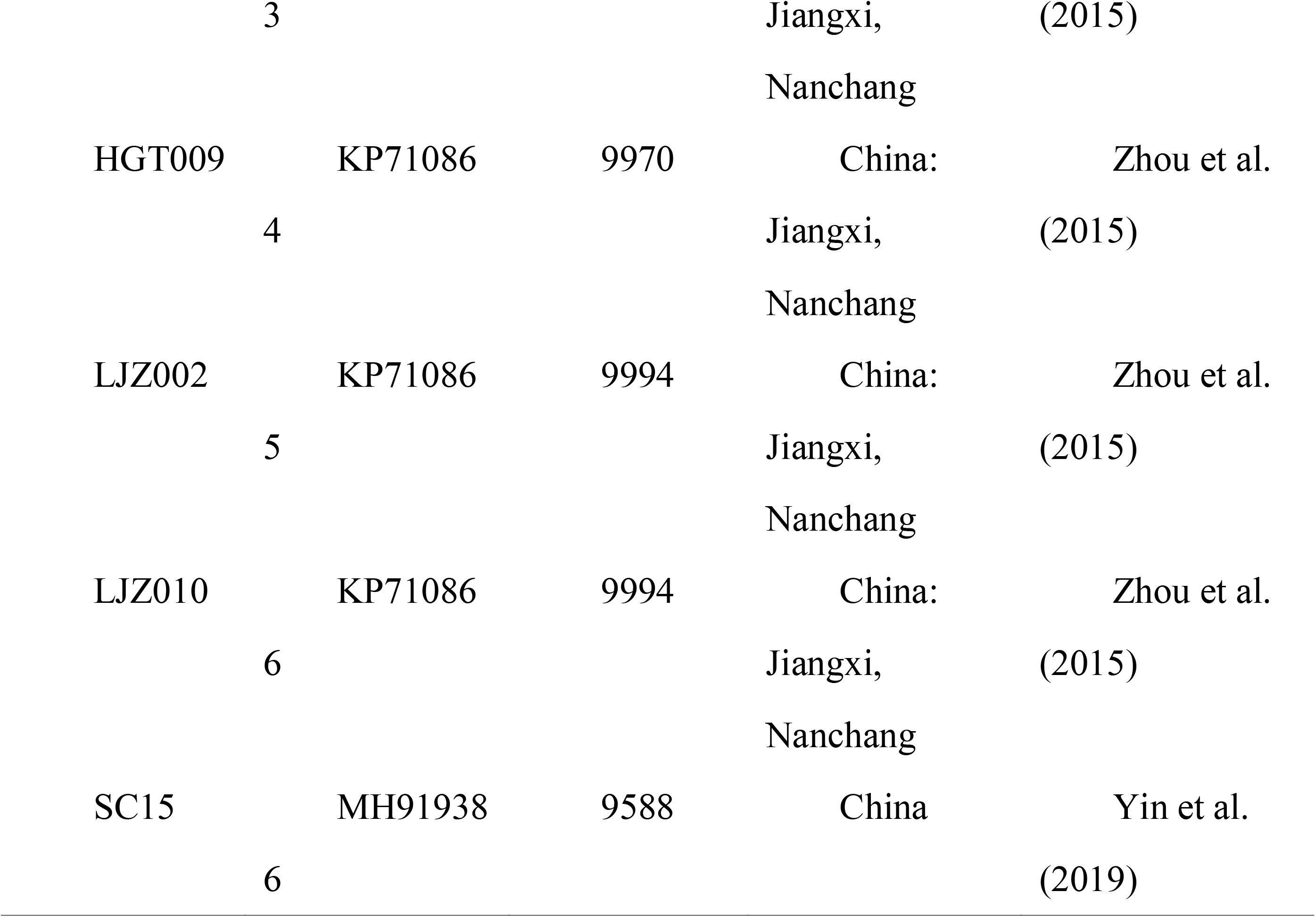
Isolates of soybean mosaic virus used in this study.

**Supplemental Figure 1.**
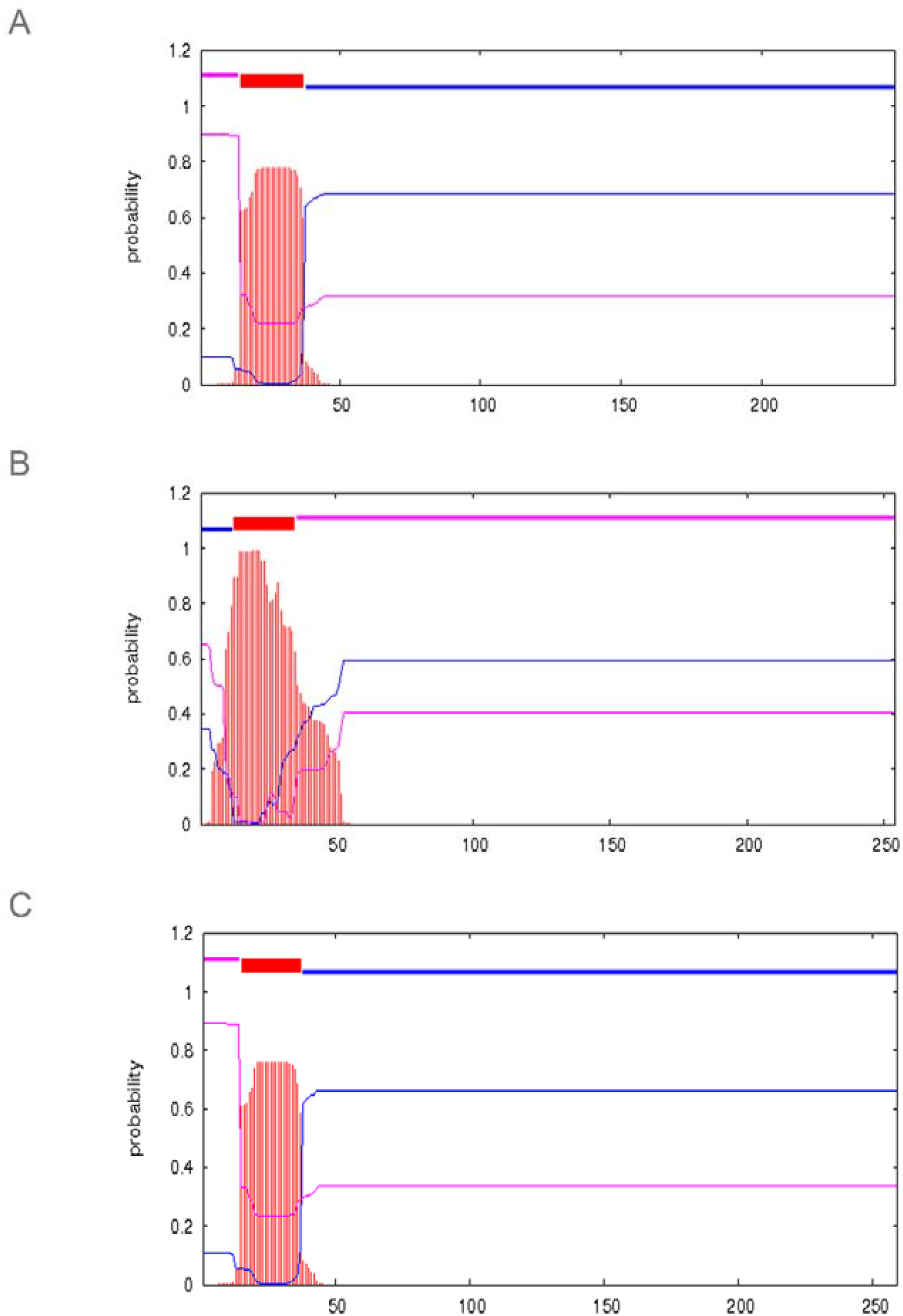
The prediction model of transmembrane regions of proteins encoded by *LOC100526921*, *LOC100812666*, and *Gm02R*. In this figure, the top line of each image represents the position of the signal sequence, the red line represents the transmembrane area, the pink line represents the signal sequence inside, and the blue line represents the signal sequence outside. A) *LOC100526921* encodes a 247 aa protein with a transmembrane helix at 15-37 aa. (B) *LOC100812666* encodes a 254 aa protein with a transmembrane helix at 13-35 aa. (C) *Rsc3K* encodes a 259 aa protein with a transmembrane helix at 15-37 aa.

**Supplemental Figure 2.**
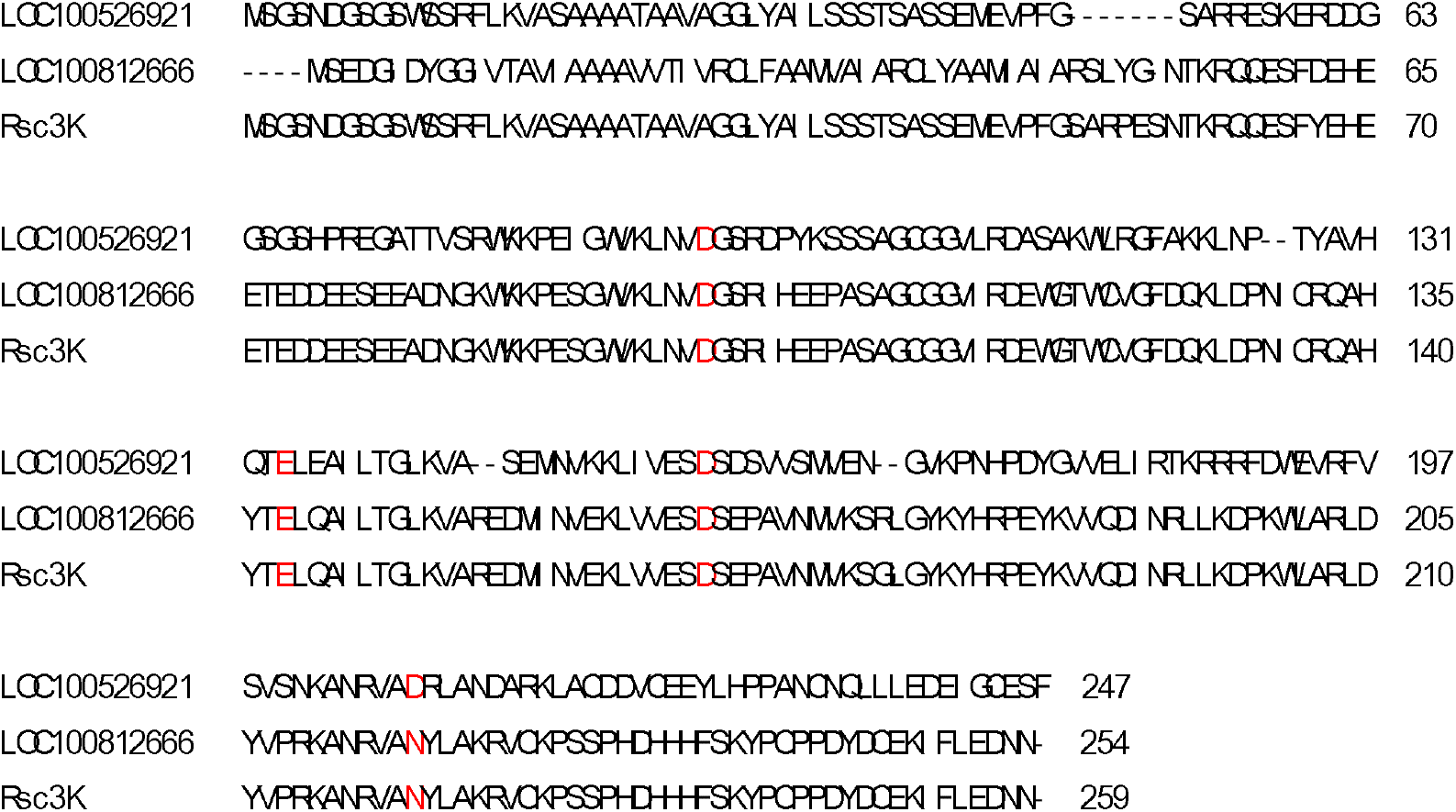
Alignment of the full-length amino acid sequences of Rsc3K, LOC100526921, and LOC100812666. The conserved DEDD model in RNase proteins has been marked with red letters. Among them, the proteins Rsc3K and LOC100812666 each contain a DEDN model (the fourth residue is asparagine).

**Supplemental Figure 3.**
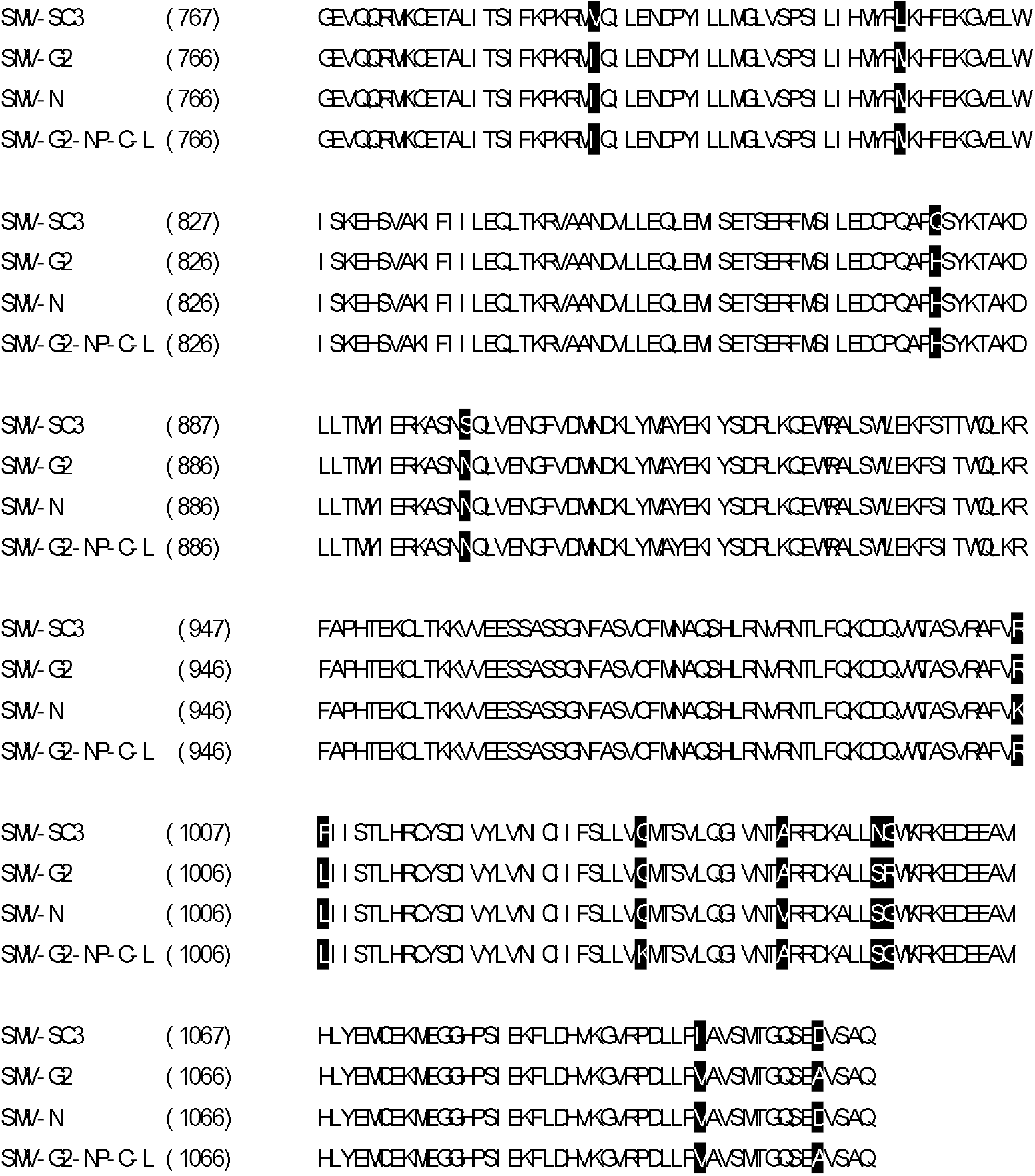
Alignment of the full-length amino acid sequences of the P3 of different SMV strains. The amino acid sequences of the P3 of SMV-SC3, SMV-G2 (NP-L), SMV-G2 (NP-C-L), and SMV-N are shown (GenBank Accession No. MH919384.1, HQ166266.1, HQ166265.1, and D00507.2 respectively). The unique amino acids of each strain are highlighted. The position of the amino acid is based on the predicted position of P3 in the polyprotein of pCB301-SC3 (Yin *et al*., 2019).

**Supplemental Figure 4.**
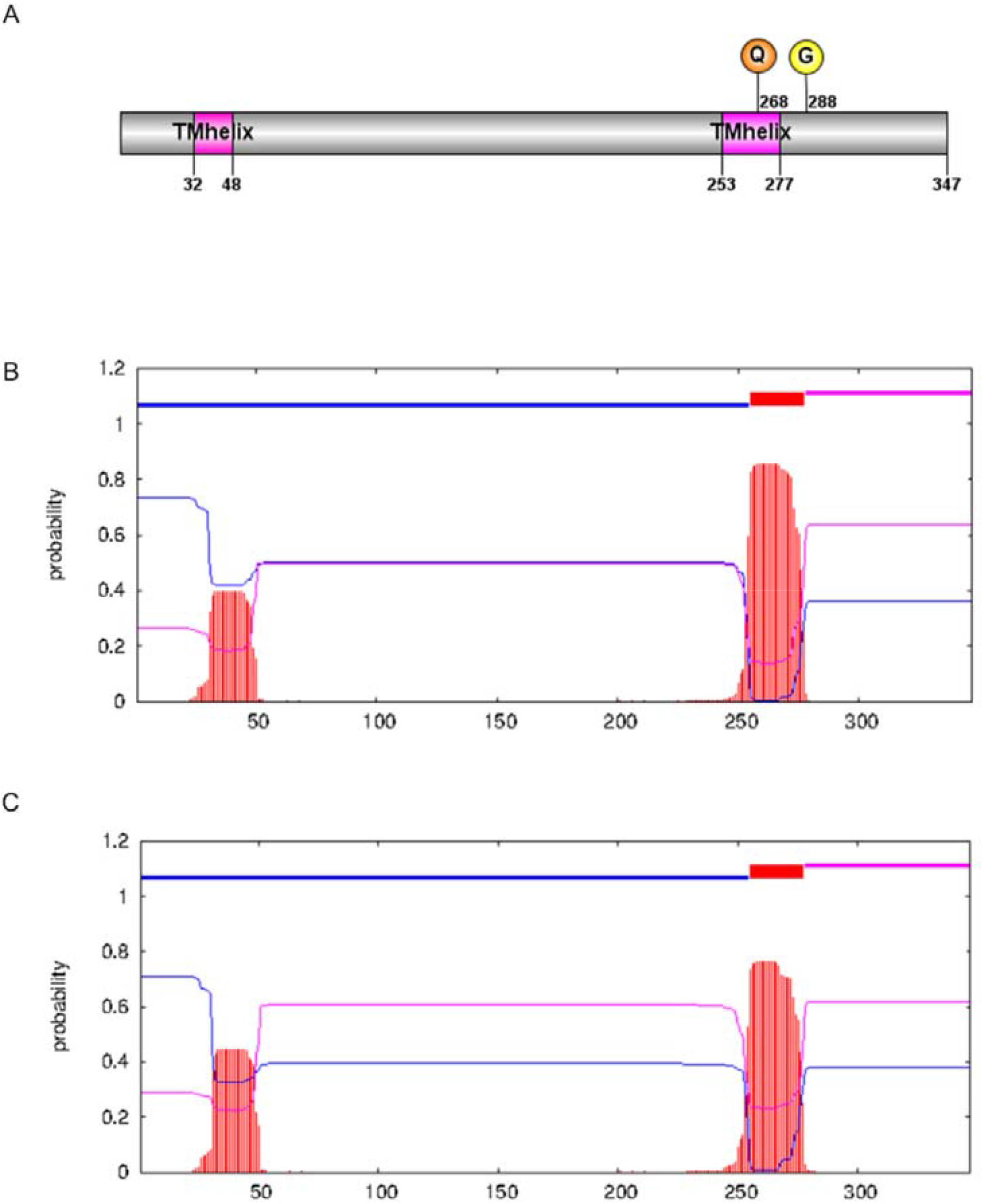
The prediction model of SMV P3 protein. The prediction model of SMV P3 by using has been drawn. (A) The SMV-SC3 P3 protein has two transmembrane helices. Among them, one virulence determining site (Q1034K) is inside the second transmembrane helix, and the other virulence determining site (G1055R) is outside the transmembrane helix region. (B) The SMV-SC3 P3 protein with two transmembrane helices at 32-48 and 253-277 aa. (C) The SMV-SC3 P3 protein (with Q1034K and G1055R variations) with two transmembrane helices at 32-48 and 253-277 aa.

**Supplemental Figure 5.**
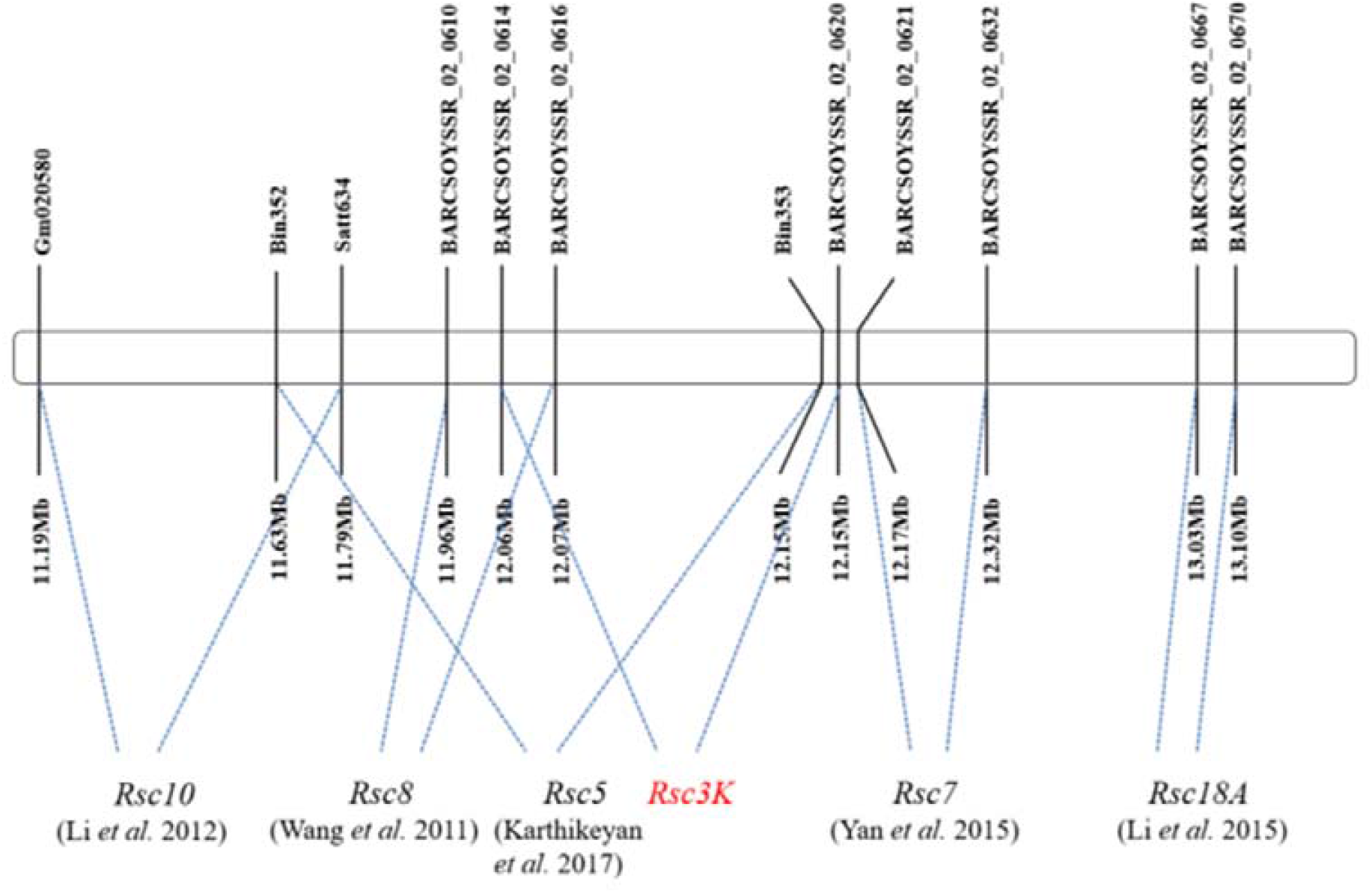
Physical maps of SMV resistance loci on chromosome 2 in Kefeng No.1. The dashed lines indicated that the mapping interval of SMV resistance loci. Among them, the *Rsc3K* locus are indicated by red letters.

Supplemental Dataset The marker genotypes of the F_2:3_ and RIL populations.

## References

Ahangaran A, Habibi MK, Mohammadi G, Winter S, García-Arenal F. 2013. Analysis of soybean mosaic virus genetic diversity in iran allows the characterization of a new mutation resulting in overcoming rsv4-resistance , Journal of General Virology 94(Pt_11) 2557–2568 ,

Chen J, Zheng HY, Lin L, Adams MJ, Antoniw JF, Zhao MF, et al. 2004. A virus related to soybean mosaic virus from pinellia ternata in china and its comparison with local soybean smv isolates. Archives of Virology, 149(2), 349–363.

Cho EK, Goodman RM. 1979. Strains of soybean mosaic virus: classification based on virulence in resistant soybean cultivars , Phytopathology 69(5) 467–470 ,

Cho EK, Goodman RM. 1982. Evaluation of resistance in soybeans to soybean mosaic virus strains1 , Crop Science 22(6) 1133–1136 ,

Chu M, Lopez-Moya JJ, Llave-Correas C, Pirone TP. 1997. Two separate regions in the genome of the tobacco etch virus contain determinants of the wilting response of tabasco pepper , Mol Plant Microbe Interact 10(4) 472–480 ,

Chowda-Reddy RV, Sun H, Hill JH, Vaino P, Wang A, Zhang Z. 2011a. Simultaneous mutations in multi-viral proteins are required for soybean mosaic virus to gain virulence on soybean genotypes carrying different r genes, Plos One 6(11) e28342,

Chowda-Reddy RV, Sun H, Chen H, Poysa V, Wang A. 2011b. Mutations in the p3 protein of soybean mosaic virus g2 isolates determine virulence on rsv4-genotype soybean, Molecular plant-microbe interactions : MPMI 24(1) 37,

Eggenberger AL, Hajimorad MR, Hill JH. 2008. Gain of virulence on rsv1-genotype soybean by an avirulent soybean mosaic virus requires concurrent mutations in both p3 and hc-pro, Molecular plant-microbe interactions : MPMI 21(7) 931,

Fu S, Yong Z, Zhi HJ, Gai JY, Yu D. 2006. Mapping of smv resistance gene rsc-7 by ssr markers in soybean, Genetica 128(1-3) 63–69,

Guo DQ, Zhi HJ, Wang YW, Gai JY, Zhou XA, Yang CL. 2005. Identification and distribution of soybean mosaic virus strains in middle and northern huang huai region of china, Chinese Journal of Oil Crop Scieves,

Hajimorad MR, Eggenberger AL, Hill JH. 2003. Evolution of soybean mosaic virus-g7 molecularly cloned genome in rsv1-genotype soybean results in emergence of a mutant capable of evading rsv1-mediated recognition, Virology 314(2) 497–509,

Hajimorad MR, Eggenberger AL, Hill JH. 2005. Loss and gain of elicitor function of soybean mosaic virus g7 provoking rsv1-mediated lethal systemic hypersensitive response maps to p3, Journal of Virology 79(2) 1215–1222,

Hajimorad MR, Eggenberger AL, Hill JH. 2006. Strain-specific p3 of soybean mosaic virus elicits rsv1-mediated extreme resistance but absence of p3 elicitor function alone is insufficient for virulence on rsv1-genotype soybean, Virology 345(1) 156–166,

Hajimorad MR, Eggenberger AL, Hill JH. 2008. Adaptation of soybean mosaic virus avirulent chimeras containing p3 sequences from virulent strains to rsv1-genotype soybeans is mediated by mutations in hc-pro, Mol Plant Microbe Interact 21(7) 937–946,

Hjulsager CK, Olsen BS, Jensen D, Cordea MI, Krath BN, Johansen IE. 2006. Multiple determinants in the coding region of pea seed-borne mosaic virus p3 are involved in virulence against sbm-2 resistance, Virology 355(1) 52–61,

Huang J, Zhang H. 2004. Development of nucleotide sequence analysis software based on windows, Bioinformatiocs,

Hull R. 2002. Matthew’s Plant Virology. New York: Academic Press

Ishibashi K, Saruta M, Shimizu T, Shu M, Kaga A. 2019. Soybean antiviral immunity conferred by dsrnase targets the viral replication complex, Nature Communications 10(1.

Johansen IE, Lund OS, Hjulsager CK, Laursen J. 2001. Recessive resistance in pisum sativum and potyvirus pathotype resolved in a gene-for-cistron correspondence between host and virus, Journal of Virology 75(14) 6609–14,

Karthikeyan A, Kai L, Hua J, Rui R, Gai JY. 2017. Inheritance fine-mapping and candidate gene analyses of resistance to soybean mosaic virus strain sc5 in soybean, Molecular Genetics and Genomics Mgg 292(4) 1–12,

Karthikeyan A, Kai L, Cui L, Yin JL, Gai JY. 2018. Fine-mapping and identifying candidate genes conferring resistance to soybean mosaic virus strain sc20 in soybean, Theoretical and Applied Genetics 131(2) 461–476,

Khatabi B, Fajolu OL, Wen RH, Hajimorad MR. 2012. Evaluation of north american isolates of soybean mosaic virus for gain of virulence on rsv-genotype soybeans with special emphasis on resistance-breaking determinants on rsv4, Molecular Plant Pathology 13(aop).

Khatabi BR, Wen H, Hajimorad M R. 2013. Fitness penalty in susceptible host is associated with virulence of soybean mosaic virus on rsv1-genotype soybean: a consequence of perturbation of hc-pro and not p3, Molecular Plant Pathology 14,

Klepadlo M, Chen P, Shi A, Mason R, Korth K, Srivastava V, Wu C. 2017. Two tightly linked genes for Soybean mosaic virus resistance insoybean, Crop Sci57:1–10

Kosambi DD. 2011. The estimation of map distances from recombination values, Annals of Human Genetics 12(1) 172–175,

Krogh A, Larsson B, von Heijne G, Sonnhammer ELL. 2001. Predicting transmembrane protein topology with a hidden markov model: application to complete genomes. Journal of Molecular Biology, 305(3), 567–580.

Kumar S, Stecher G, Tamura K. 2016. Molecular evolutionary genetics analysis version 7.0 for bigger datasets. Moleculer. Molecular Biology and Evolution, 33, 1870–1874.

Li C, Yang YQ, Wang DG, Li H, Zheng GJ, Wang T, Zhi HJ. 2012. Studies on mapping and inheritance of resistance genes to smv strain sc10 in soybean, Scientia Agricultura Sinica,Scientia Agricultura Sinica 45(21): 4335–4342,

Li K, Yang QH, Zhi HJ, Gai JY. 2010. Identification and distribution of soybean mosaic virus strains in southern china, Plant Disease 94(3) 351–357,

Li K, Ren R, Adhimoolam K, Gao L, YuanYQ, Liu, Z, et al. 2015. Genetic analysis and identification of two soybean mosaic virus resistance genes in soybean [glycine max (l.) merr]. Plant Breeding.

Ma Y, Li H, Wang DG, Li N, Zhi HJ. 2010. Molecular mapping and marker assisted selection of soybean mosaic virus resistance gene rsc12 in soybean, Legume Genomics and Genetics 1,

Ma Y, Wang DG, Li HC, et al. 2011. Fine mapping of the rsc14q locus for resistance to soybean mosaic virus in soybean, Euphytica 181(1) 127–135,

Matthews, Ef R. 2002. Matthews’ plant virology, Matthews Plant Virology,

Maroof M, Tucker DM, Tolin SA. 2008. Genomics of Viral–Soybean Interactions, Springer New York,

Mohammad A, Sher H. 2002. Evaluation of resistance in soybean germplasm to soybean mosaic potyvirus under field conditions, Journal of Biological Sciences 2(9) 601–604,

Nakagawa T, Kurose T, Hino T, Tanaka K, Kimura T. 2007. Development of series of gateway binary vectors pgwbs for realizing efficient construction of fusion genes for plant transformation, Journal of Bioscience and Bioengineering 104(1) 34–41,

Nowotny M, Gaidamakov SA, Crouch RJ, Yang W. 2005. Crystal structures of RNase H bound to an RNA/DNA hybrid: substrate specificity and metaldependent catalysis. Cell 121(7) 1005–1016.

Ooijen J, Van JW. 2006. JoinMap 4 Software for the calculation of genetic linkage maps in experimental populations,

Padgett HS, Watanabe Y, Beachy RN. 1997. Identification of the tmv replicase sequence that activates the n gene-mediated hypersensitive response, Mol, Plant-Microbe Interact,10(6) 709–715,

Ren R, Liu SC, Karthikeyan A, Wang T, Niu HP, Yin JL, et al. 2017. Fine-mapping and identification of a novel locus rsc15 underlying soybean resistance to soybean mosaic virus, Theoretical and Applied Genetics,

Salvador B, Delgadillo MO, Saenz P, Garcia JA, Simonmateo C. 2008. Identification of plum pox virus pathogenicity determinants in herbaceous and woody hosts, Molecular plant-microbe interactions : MPMI 21(1) 20,

Seo JK, Lee SH, Kim KH. 2009. Strain-specific cylindrical inclusion protein of soybean mosaic virus elicits extreme resistance and a lethal systemic hypersensitive response in two resistant soybean cultivars, Molecular plant-microbe interactions : MPMI 22(9) 1151,

Tusnády GE and Simon I. 1998. Principles governing amino acid composition of integral membrane proteins: application to topology prediction. Journal of Molecular Biology, 283(2), 489–506.

Tusnády GE and Simon I. 2001. The hmmtop transmembrane topology prediction server. Bioinformatics(9), 849–850.

Voorrips R. 2002. MapChart: software for the graphical presentation of linkage maps and QTLs, J Hered 93:77–78

Wang DG, Ying M, Ning L, Yang Z, Zheng GJ, Zhi HJ. 2011a. Fine mapping and identification of the soybean Rsc4 resistance candidate gene to soybean mosaic virus, Plant Breeding 130(6.

Wang DG, Ying M, Yang YQ, Ning L, Li C, Song YP, et al. 2011b. Fine mapping and analyses of Rsc8 resistance candidate genes to soybean mosaic virus in soybean, Theoretical and Applied Genetics 122(3) p,555–565,

Wang DG, Li K, Zhi HJ. 2018. Progresses of resistance on soybean mosaic virus in soybean, Scientia Agricultura Sinica,

Wrather JA, Anderson TR, Arsyad DM, Gai J, Yorinori JT. 1997. Soybean disease loss estimates for the top 10 soybean producing countries in 1994, Plant Disease 81(1) 107–110,

Wang X, Gai J, Pu Z. 2003. Classification and distribution of strain groups of soybean mosaic virus in middle and lower huang-huai and changjiang valleys, Soybean Science,

Wang Y, Khatabi B, Hajimorad MR. 2015. Amino acid substitution in p3 of soybean mosaic virus to convert avirulence to virulence on rsv4-genotype soybean is influenced by the genetic composition of p3, Molecular Plant Pathology 16(3).

Wang YW, Zhi HJ, Guo DQ, Gai JY, Li HC. 2005. Classification and distribution of strain groups of soybean mosaic virus in northern china spring planting soybean region, Soybean Science,

Wen RH, Khatabi B, Ashfield T, Maroof M, Hajimorad MR. 2013. The hc-pro and p3 cistrons of an avirulent soybean mosaic virus are recognized by different resistance genes at the complex rsv1 locus, Molecular Plant-microbe Interactions:MPMI 26(2.

Yan H, Wang H, Cheng H, Hu Z, Chu S, Zhang G, et al. 2015. Detection and fine-mapping of sc7 resistance genes via linkage and association analysis in soybean, Journal of Integrative Plant Biology 57(008) 722–729,

Yang QH, Gai JY. 2011. Identification inheritance and gene mapping of resistance to a virulent soybean mosaic virus strain sc15 in soybean, Plant Breeding 130(2) 128–132,

Yang XF, Yang YQ, Zheng GJ, Zhi HJ, Li XH. 2013. Fine mapping of resistance genes to smv strains sc6 and sc17 in soybean, ACTA AGRONOMICA SINICA 39(2) 216,

Yang Y, Zheng G, Han L, Wang D. 2013. Genetic analysis and mapping of genes for resistance to multiple strains of soybean mosaic virus in a single resistant soybean accession pi 96983, Theoretical and Applied Genetics 126(7) 1783–1791,

Yin JL, Liu H, Xiang WY, Jin TT, Zhi HJ. 2019. Discovery of the agrobacterium growth inhibition sequence in virus and its application to recombinant clone screening, AMB Express 9(1.

Yin JL, Wang LQ, Jin TT, Nie Y, Liu H, et al. 2021. A cell wall-localized NLR confers resistance to Soybean mosaic virus by recognizing viral-encoded cylindrical inclusion protein, Molecular Plant 1674–2052

Z Divéki, K Salánki, and E Balázs. 2004. The necrotic pathotype of the cucumber mosaic virus cmv) ns strain is solely determined by amino acid 461 of the 1a protein, Molecular Plant-Microbe Interactions 17(8) 837–845,

Zhang C, Hajimorad MR, Eggenberger AL, Tsang S, Whitham SA, Hill JH. 2009. Cytoplasmic inclusion cistron of soybean mosaic virus serves as a virulence determinant on rsv3-genotype soybean and a symptom determinant, VIROLOGY -NEW YORK-391(2.

Zhang C, Bradshaw JD, Whitham SA, Hill JH. 2010. The development of an efficient multipurpose bean pod mottle virus viral vector set for foreign gene expression and rna silencing, Plant Physiology 153(1) 52–65,

Zhang HY, Cui XY, Chen X, Zhi H, Zhang S, Zhao L. 2012. Determination of the complete genomic sequence and molecular biological analysis of soybean mosaic virus, Canadian Journal of Plant Pathology 34(2) 1–10,

Zheng GJ, Yang YQ, MA Y, Yang XF, Chen SY, Ren R, et al. 2014. Fine mapping and candidate gene analysis of resistance gene rsc3q to soybean mosaic virus in qihuang 1, Journal of Integrative Agriculture,

Zhang J, H. 2009. Control technology of soybean mosaic virus disease, China Science and Technology Fortune Magazine 06:74

Zhang ZY, Chen SY, GAI JY. 1999. Molecular markers linked to rsa resistant to soybean mosaic virus. Chinese Science Bulletin.

Zhi HJ, Yuan Y, Gao L, Liu ZT, LI K, Zhong YK, Ren R, Karthikeyan A, et al. 2016. Genetic analysis and identification of two soybean mosaic virus resistance genes in soybean [glycine max l,) merr], Plant Breeding 134(6) 684–695,

Zhou GC, Shao ZQ, Ma FF, Wu P, Wu XY, Xie ZY, et al. 2015. The evolution of soybean mosaic virus: an updated analysis by obtaining 18 new genomic sequences of chinese strains/isolates. Virus Research, 208, 189–198.

Zong TX, Yin JL, Jin TT, Wang LQ, Luo MX, Li K, et al. 2020. A dnaj protein that interacts with soybean mosaic virus coat protein serves as a key susceptibility factor for viral infection - sciencedirect, Virus Research 281,

